# Zinc-Enhanced Activity of an Antimicrobial Halogenated Phenazine Against *Streptococcus mutans* and Other Gram-positive Bacteria

**DOI:** 10.1101/2025.07.10.664208

**Authors:** Jessica K. Kajfasz, Hannah B. Hosay, Qiwen Gao, Robert W. Huigens, José A. Lemos

## Abstract

Halogenated phenazine (HP) compounds have shown promise as antimicrobial agents, particularly against biofilm-associated Gram-positive pathogens. Among these compounds, HP-29 demonstrates potent activity against methicillin-resistant *Staphylococcus aureus* by inducing rapid iron starvation. As maintenance of trace metals homeostasis is critical for the survival of *Streptococcus mutans*, this study investigated the antimicrobial efficacy of HP-29 and the impact of metal supplementation on this major oral and occasional systemic pathogen. As anticipated, HP-29 inhibited *S. mutans* growth in a dose-dependent manner, with iron supplementation alleviating the antimicrobial effect. Cobalt, manganese, or nickel supplementation also mitigated the inhibitory activity of HP-29 but, unexpectedly, the addition of zinc greatly enhanced HP-29 antimicrobial activity. This zinc-driven potentiation of HP-29 extended to other Gram-positive pathogens, including *Enterococcus faecalis* and *S. aureus*. Inductively coupled plasma mass spectrometry analysis revealed that intracellular iron content decreased significantly following exposure to HP-29. At the same time, exposure to HP-29 led to a slight increase in intracellular zinc, mirroring the increase observed in cells exposed to excess zinc. When combined with zinc, HP-29 triggered a 5-fold increase in intracellular zinc and reduced manganese levels by ∼50%. Transcriptome analysis showed that HP-29, with or without zinc, altered expression of genes linked to iron and manganese uptake as well as zinc efflux, suggesting broad disruption of metal ion regulation. These findings highlight HP-29 as a potent antimicrobial that broadly impairs metal homeostasis. The unexpected synergy of HP-29 with zinc points toward a promising dual-agent therapeutic strategy against Gram-positive pathogens.

**IMPORTANCE:** Widespread development of antibiotic resistance has created a constantly moving target when combating infectious microbes. Here, we further explore an antimicrobial halogenated phenazine, HP-29, which is effective against Gram-positive bacteria through disruption of intracellular trace metal equilibrium. We showed that HP-29 inhibits growth of the oral and systemic pathogen *Streptococcus mutans* and that its antimicrobial effect is greatly potentiated by the addition of zinc. The zinc-mediated enhancement of HP-29’s efficacy was also observed in other Gram-positive pathogens, including *Enterococcus faecalis* and *Staphylococcus aureus*. Intracellular trace metal quantifications and transcriptome analysis confirmed that HP-29 treatment impairs trace metal homeostasis, an outcome that is exacerbated when *S. mutans* is treated with both HP-29 and zinc. The observed synergy of HP-29 with zinc supports the development of a dual-agent therapeutic strategy against Gram-positive pathogens.

## INTRODUCTION

The constant and fast emergence of multidrug (MDR) resistant bacteria is a topic of great concern, prompting efforts to develop new antibacterial strategies (1, 2). Recent work has shown that certain halogenated phenazine (HP) analogs can be highly effective against Gram-positive pathogens, including those with broad resistance to traditional antibiotics (3). This series of compounds was inspired by natural bacterial competition, where strains of *Pseudomonas aeruginosa* were found to secrete pyocyanin, a phenazine antibiotic and virulence factor, that eliminated *Staphylococcus aureus* from the lungs of cystic fibrosis patients (4). Focused libraries of HPs have been synthesized and found to effectively kill Gram-positive pathogens, both in the planktonic and the biofilm state (5–10). Supporting previous observations that HP compounds bind divalent metal cations (6–10), transcriptional profiling of methicillin-resistant *S. aureus* (MRSA) biofilms following treatment with HP-14 (halogenated phenazine analog 14) revealed a rapid induction in expression of gene clusters associated with iron uptake (11). Continued efforts to explore new HP analogs through chemical synthesis and microbiological studies led to the identification of HP-29, which was highly effective against MRSA and *Enterococcus faecalis* among other major Gram-positive pathogens (10). When provided as a topically applied ointment, HP-29 treatment significantly reduced bacterial burden on wounds of mice infected with either *E. faecalis* or *S. aureus* (10).

*Streptococcus mutans* is a keystone pathogen of dental caries and one of the causative agents of infective endocarditis, a life-threatening infection of heart valve endothelium (12–15). As a member of the oral biofilm community, the *S. mutans* lifestyle demands the ability to adapt to large fluctuations in the availability and content of nutrients, which derive almost exclusively from the human host diet (16). Transition metals are essential micronutrients for all domains of life, as the function of about 40% of all enzymes is dependent upon a metal cofactor (17–19). Studies conducted by our group and others have shown that the ability to maintain trace metal homeostasis, achieved by proteins dedicated to the sensing, import, and efflux of metals, is a critical aspect of *S. mutans* pathophysiology (20–27). Several transport systems responsible for the import and efflux of trace metals in *S. mutans* have been characterized, including those for iron (*feoABC*, *sloABC*, *smu.995-998*) (20, 26), manganese (*sloABC*, *mntH*, *mntE*) (25, 26, 28), zinc (*adcABC*, *zccE*) (21, 22), and copper (*copA*) (27). As over-accumulation of metals is associated with toxicity, *S. mutans* relies on metal-sensing regulators (metalloregulators) to tightly govern metal uptake and efflux, including SloR (iron and manganese uptake), AdcR (zinc uptake), CopY (copper export), and ZccR (zinc export) (21, 22, 29–32). Additionally, the peroxide sensor PerR, a member of the Fur (ferric uptake regulator) protein family, requires a metal cofactor and has been also linked to expression of genes associated with the transport and storage of iron and manganese (33, 34).

In this investigation, we assessed the potential of HP-29 to serve as an antimicrobial agent against *S. mutans*. Growth of *S. mutans* was inhibited by HP-29 in a dose-dependent manner that was alleviated by supplementation with iron. Expanding the study to include other trace metals revealed that cobalt, manganese and nickel could also alleviate HP-29 inhibitory activity. However, zinc supplementation greatly enhanced the antimicrobial efficacy of HP-29, an observation that was extended to other Gram-positive bacteria. Intracellular trace metal quantifications and transcriptome analysis of *S. mutans* cultures treated with HP-29 alone or combined with a non-inhibitory concentration of zinc revealed that HP-29 broadly disrupts trace metal homeostasis, and that this effect is further exacerbated upon the addition of zinc. This study confirms HP-29 as a potent antimicrobial agent against Gram-positive pathogens that disrupts intracellular metal homeostasis while also revealing the therapeutic potential of combining HPs with zinc to treat bacterial infections.

## RESULTS

### HP-29 is inhibitory to oral streptococci in a metal-dependent manner

To explore the antimicrobial potential of HP-29 for the prevention and treatment of *S. mutans* infections, we first tested the ability of *S. mutans* UA159 to grow in the presence of HP-29. Growth curves revealed a dose-dependent effect with 1.5 μM HP-29 completely inhibiting its growth, with similar inhibition seen for *S. sanguinis* and *S. gordonii* (**Fig. 1A**). MIC determinations performed using the broth microdilution method confirmed the efficacy of HP-29 against these oral streptococci, with *S. sanguinis* showing the greatest sensitivity, with an MIC of 0.125 μM and *S. mutans* showing the highest tolerance, with an MIC of 0.5 μM (**Fig. 1B**, **Table 1**). The chemical structure of HP-29 is shown in **Figure 1C**.

**Fig. 1.**
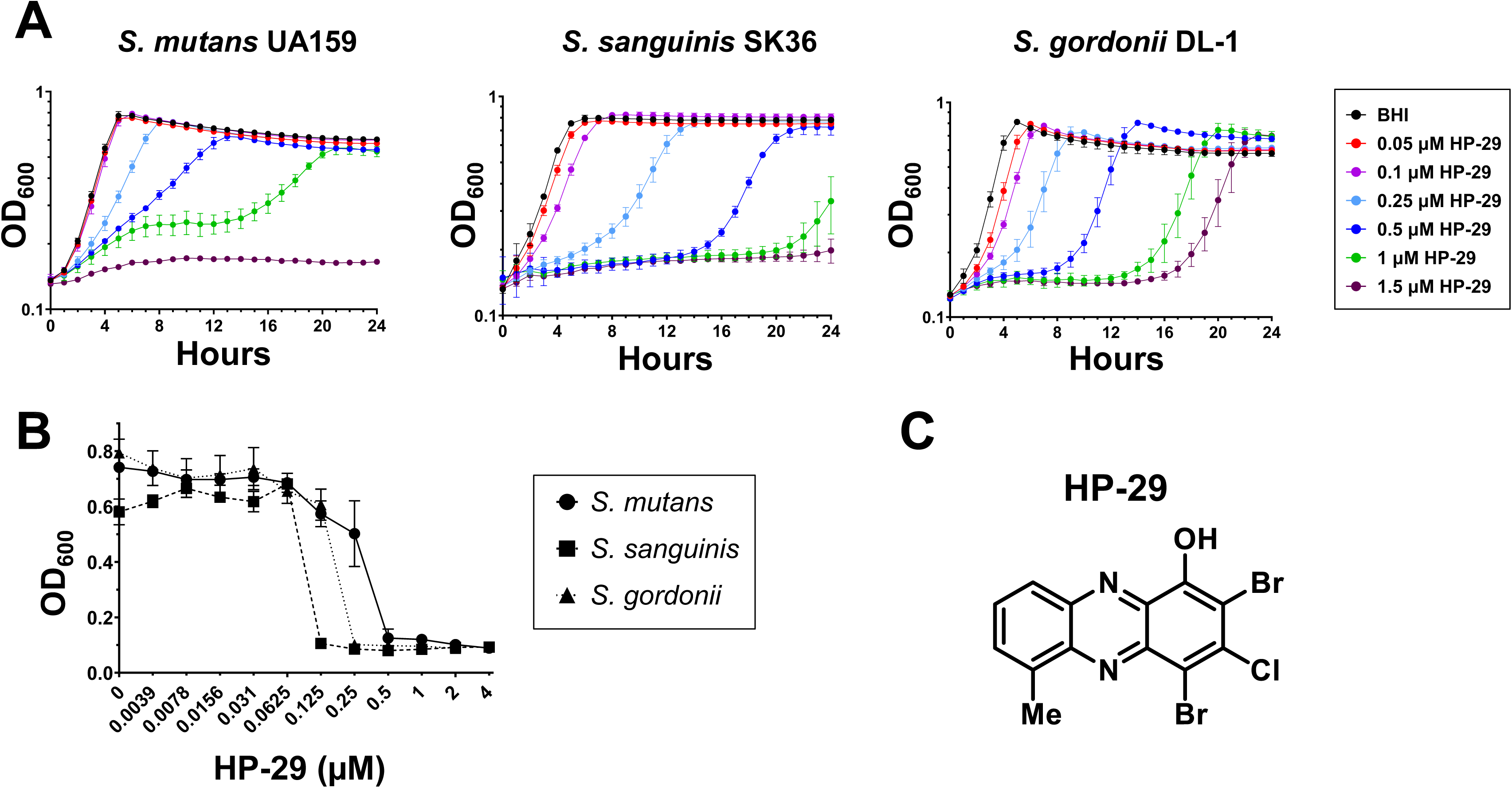
The novel compound HP-29 inhibits growth and survival of oral streptococci. The oral streptococcus strains *S. mutans* UA159, *S. sanguinis* SK36, and *S. gordonii* DL-1 were grown in BHI medium and exposed to the halogenated phenazine HP-29 in (A) growth curve assays or (B) minimal inhibitory concentration (MIC) assays. (C) Chemical structure of HP-29. Data represent averages and standard deviations of at least 3 independent experiments.

**Table 1.**
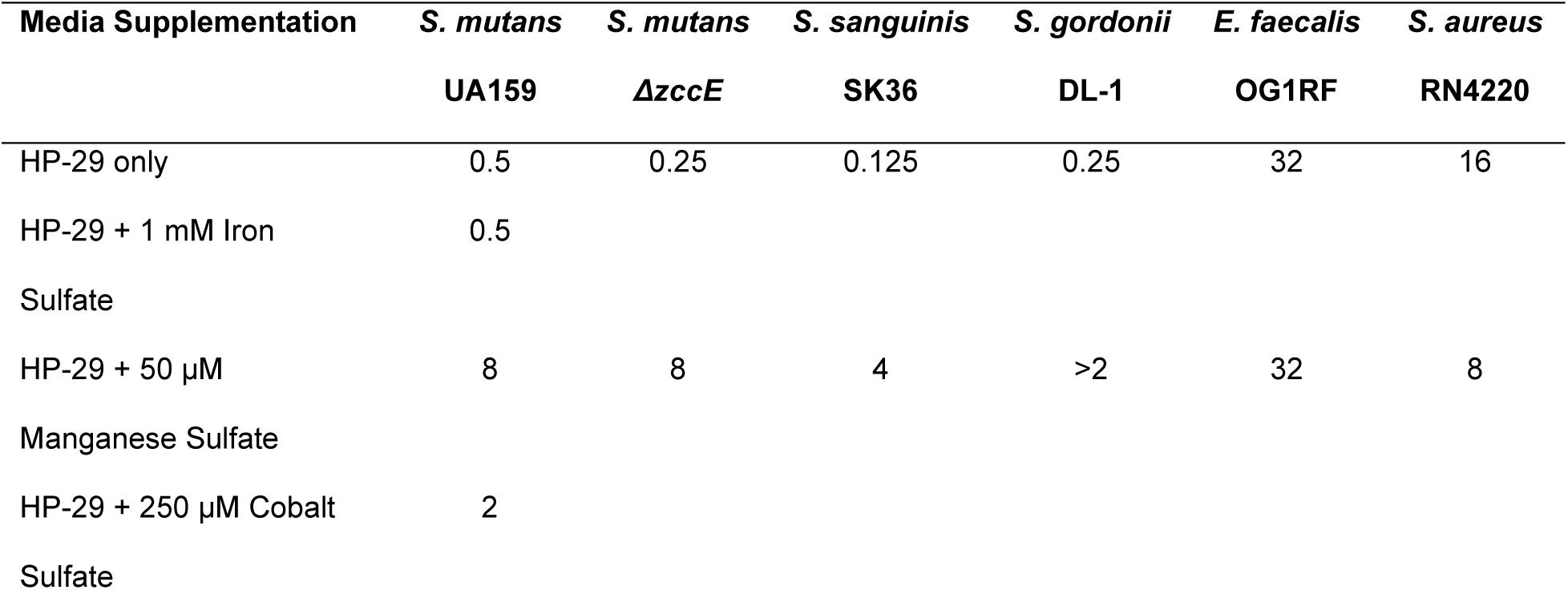
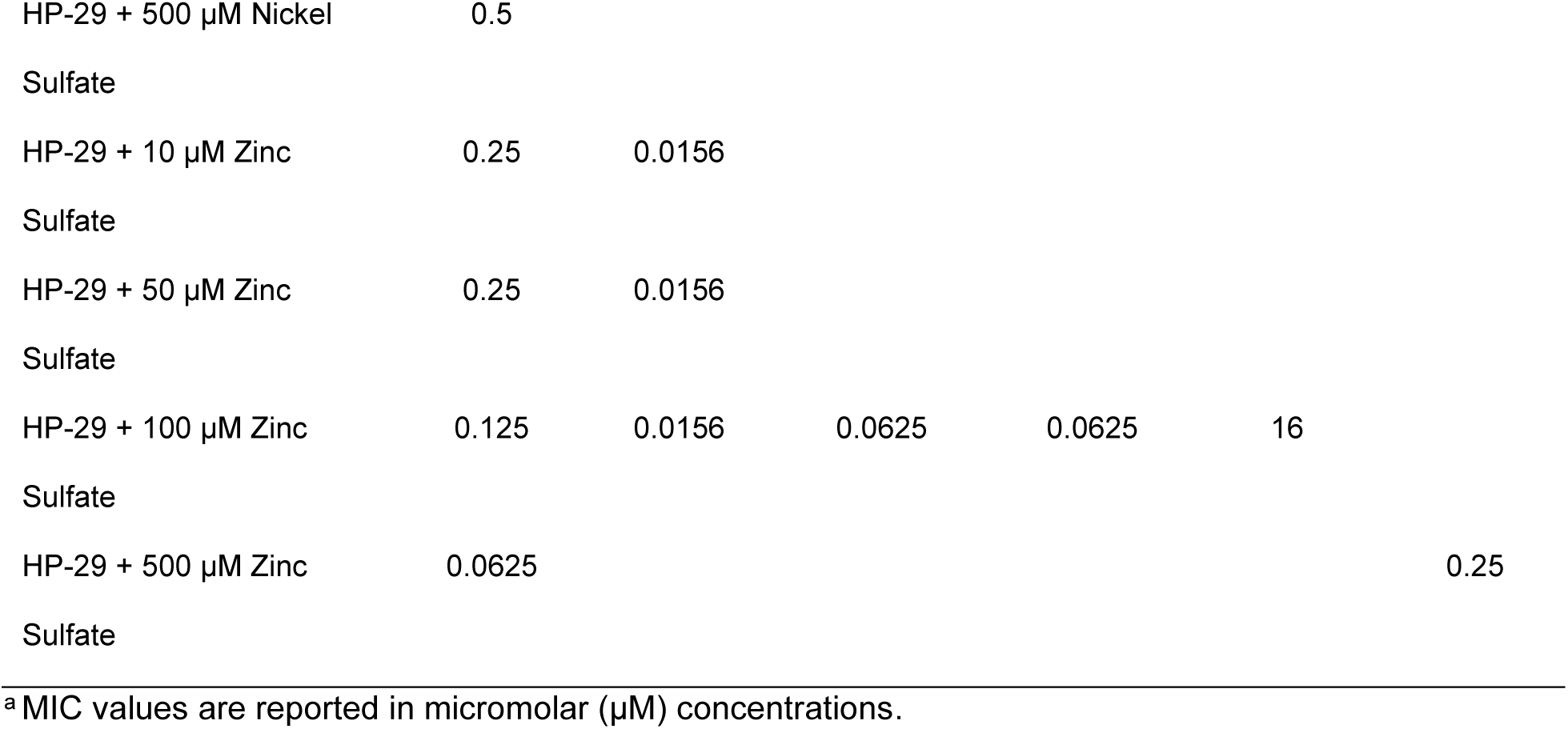
Summary of MIC values^a^ for HP-29 with or without supplementation with metals.

Previously, we have shown that brain-heart infusion (BHI) medium, the media used in the growth curve and MIC assays, contains low concentrations of the essential trace metals iron (∼ 6 μM), zinc (∼11 μM), and manganese (<1 μM) (25). Knowing that HP-29 triggers rapid iron starvation in *S. aureus* (10), we repeated the HP-29 growth kinetic assay in BHI supplemented with a sub-inhibitory concentration of iron, focusing first on the response of *S. mutans*. As expected, the addition of 1 mM FeSO_4_, the highest feasible concentration due to the tendency of iron to precipitate, partially rescued the growth inhibitory effect of HP-29 (**Fig. 2A**). Given that HP-29 can bind other metals *in vitro* (3, 10), we next sought to evaluate the effect of supplementation with sub-inhibitory concentrations of other divalent metals on HP-29 activity. Similar to iron, supplementation with manganese, nickel, or cobalt alleviated the inhibitory activity of HP-29 (**Fig. 2B-D**). Notably, while the addition of iron, nickel, or cobalt offered partial rescue of *S. mutans* UA159 growth, the addition of manganese resulted in a complete growth restoration. Unexpectedly, zinc supplementation caused the opposite effect, strongly potentiating the growth inhibitory effects of HP-29 (**Fig. 2E**).

**Fig. 2.**
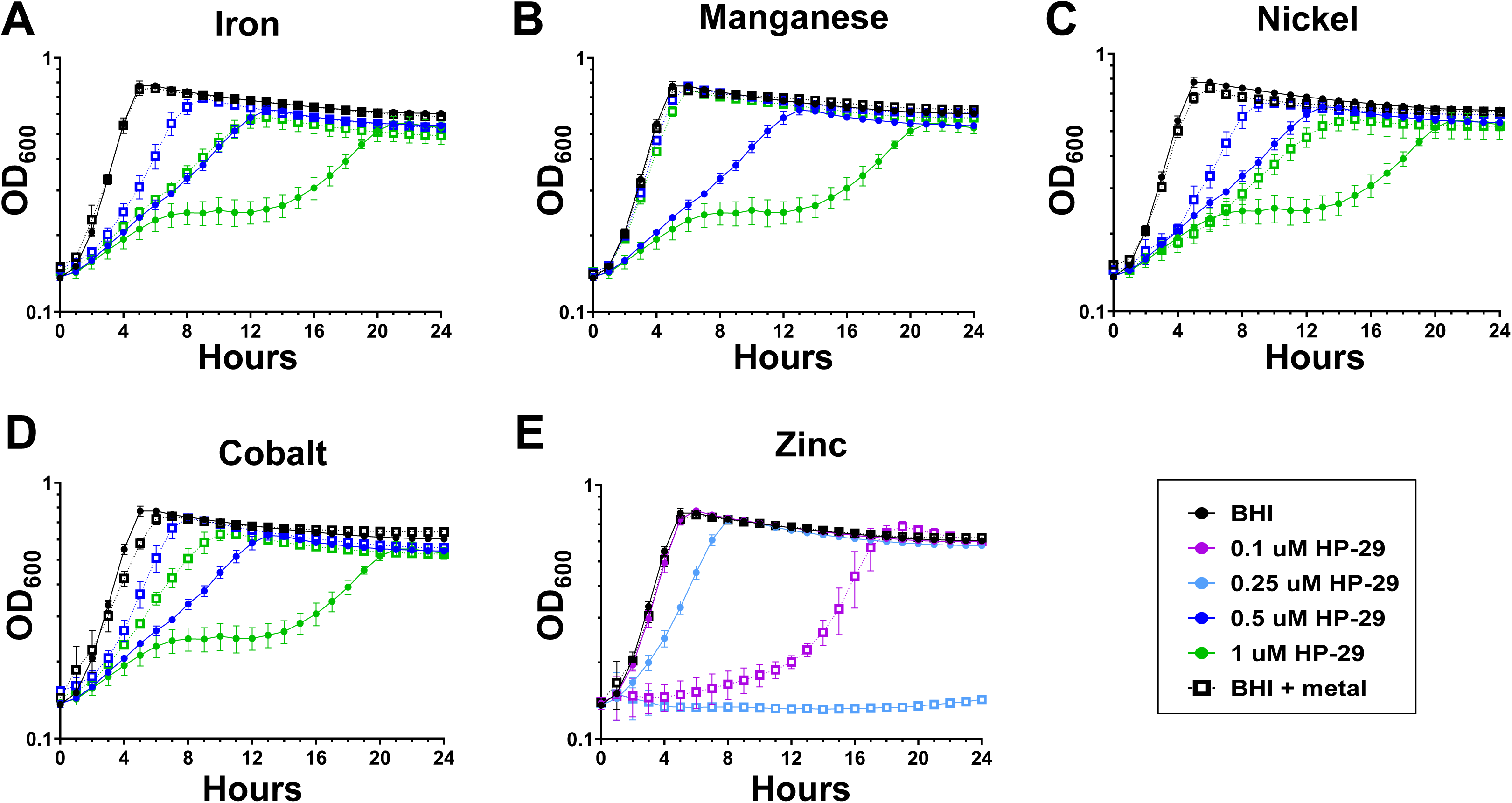
Divalent metal cations can either rescue or exacerbate the ability of HP-29 to inhibit growth of *S. mutans*. *S. mutans* UA159 was grown in BHI media containing 0.1 – 1 μM HP-29. The media was supplemented with the divalent metal cations (A) iron, 1 mM, (B) manganese, 0.5 mM, (C) nickel, 0.5 mM, (D) cobalt, 0.25 mM, or (E) zinc, 0.5 mM. Closed circles represent growth in the presence of HP-29, while open squares of the same color represent the addition of metal. Each panel includes two representative concentrations of HP-29. Data represent averages and standard deviations of at least 3 independent experiments.

To confirm these results, we assessed the MIC of HP-29 when combined with each individual metal (**Fig. 3A**, **Table 1**). Supporting the trends observed in the growth curves, cobalt or manganese supplementation raised the HP-29 MIC from 0.5 μM to 2 μM or 8 μM, respectively. Despite the effects noted on growth curve assays, nickel or iron supplementation did not affect the HP-29 MIC. Most notably, the addition of 0.5 mM ZnSO_4_ lowered HP-29 MIC to 0.0625 μM HP-29, an 8-fold increase in sensitivity as compared to exposure to HP-29 alone (**Table 1**). Since the addition of zinc resulted in such a strong phenotype, a zinc titration was performed revealing that as little as 0.01 mM ZnSO_4_ increased *S. mutans* sensitivity to HP-29 by 2-fold (**Fig. 3B**, **Table 1**). Previously, we showed that the zinc exporter ZccE mediates the high zinc tolerance of *S. mutans* (21). Here, we show that the Δ*zccE* strain was slightly more sensitive to HP-29 than the parent strain in BHI (MIC of 0.25 µM for Δ*zccE* as compared to 0.5 µM for UA159). As expected, the combination of HP-29 with sub-inhibitory concentrations of ZnSO_4_ was highly inhibitory to the growth of the Δ*zccE* strain; for instance, in the presence 0.05 mM zinc, the MIC of HP-29 decreased 16-fold to 0.0156 μM (**Fig. 3C, Table 1).** As seen with the parent strain, manganese supplementation was highly effective at rescuing sensitivity of Δ*zccE* to HP-29, increasing the MIC by 32-fold (**Fig. 3C**, **Table 1**).

**Fig. 3.**
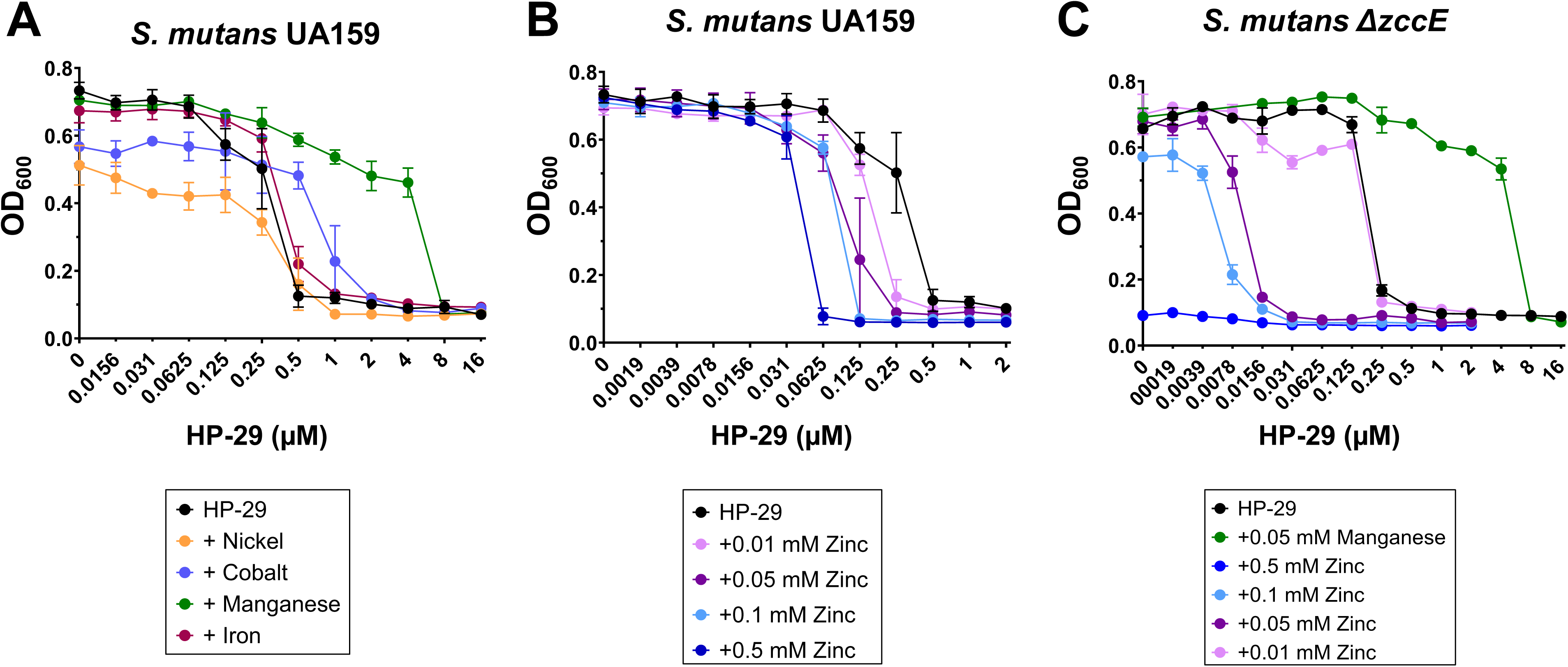
Divalent metal cations can either rescue or potentiate the MIC of HP-29 against *S. mutans*. (A) HP-29 MIC assay for *S. mutans* UA159 with the addition of trace metals: 0.5 mM nickel, 0.25 mM cobalt, 0.05 mM manganese, 1 mM iron. (B) A titration assay shows that as little as 0.01 mM zinc increases sensitivity of *S. mutans* UA159 to HP-29. (C) The HP-29 MIC assays with supplementation of manganese or zinc were repeated with the *S. mutans ΔzccE* strain. Data represent averages and standard deviations of at least 3 independent experiments.

Knowing that metal tolerance varies widely among bacteria (35, 36), the ability of divalent metal cations to rescue or potentiate the inhibitory effect of HP-29 in other Gram-positive organisms was evaluated. Here, we used only manganese or zinc as representatives of metals that either rescue or potentiate the antimicrobial activity of HP-29 in *S. mutans*. Like in *S. mutans*, manganese supplementation greatly enhanced the HP-29 MIC for *S. sanguinis* and *S. gordonii*, while zinc supplementation increased sensitivity by 2– and 4-fold, respectively (**Fig. 4A-B**). We also assessed the inverse correlation of manganese and zinc when combined with HP-29 against *E. faecalis* and *S. aureus*, organisms of particular medical concern to humans due to their association with antibiotic resistance (37–40). While manganese supplementation failed to alleviate the HP-29 sensitivity in *E. faecalis*, the addition of zinc increased its HP-29 MIC by 2-fold (from 32 μM to 16 μM) (**Fig. 4C**). In S. *aureus*, the addition of zinc caused a remarkable 64-fold increase in HP-29 sensitivity (from 16 μM to 0.25 μM) while manganese supplementation lowered the MIC from 16 to 8 μM (**Fig. 4D**).

**Fig. 4.**
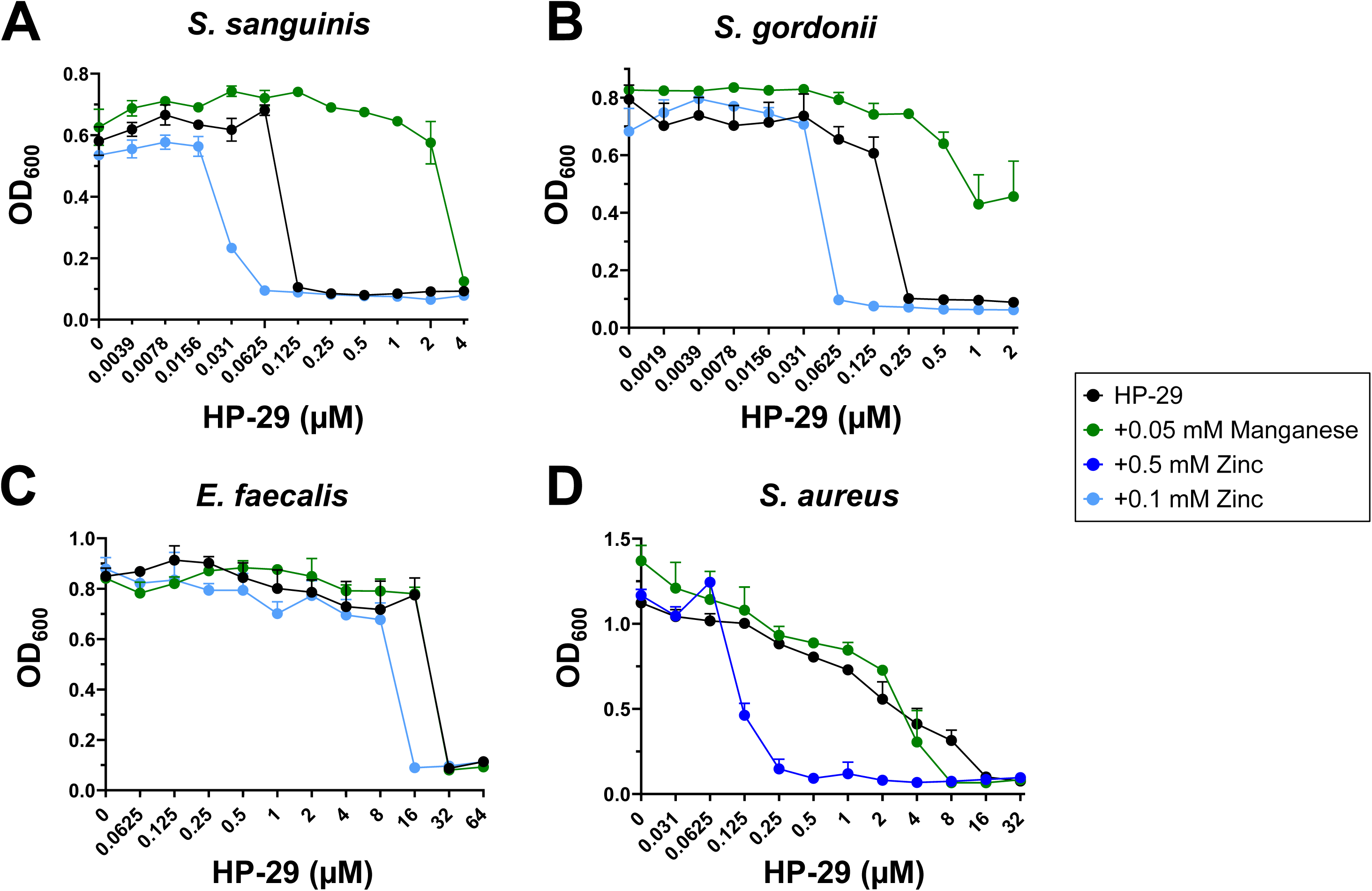
Divalent metal cations can either rescue or exacerbate the MIC of HP-29 against a panel of bacterial strains. MIC assays were performed in the presence of manganese or zinc to test for the ability to rescue or potentiate the sensitivity to HP-29 for (A) *S. sanguinis* (B) *S. gordonii*, (C) *E. faecalis*, (D) *S. aureus*. Data represent averages and standard deviations of at least 3 independent experiments.

In previous work, HP-29 has shown rapid binding to divalent iron in UV-vis spectroscopy experiments (10). Based on the rescue and potentiation profiles observed in the presence of different metals, we investigated the ability of HP-29 to bind other divalent metal cations. UV-vis spectroscopy revealed that the binding capacity of HP-29 extends to a range of divalent metal cations, including nickel, cobalt, manganese, magnesium, and zinc (**Fig. 5A**) with HP-29 binding to these metal cations in a 2:1 ratio (**Fig. 5B**).

**Fig. 5.**
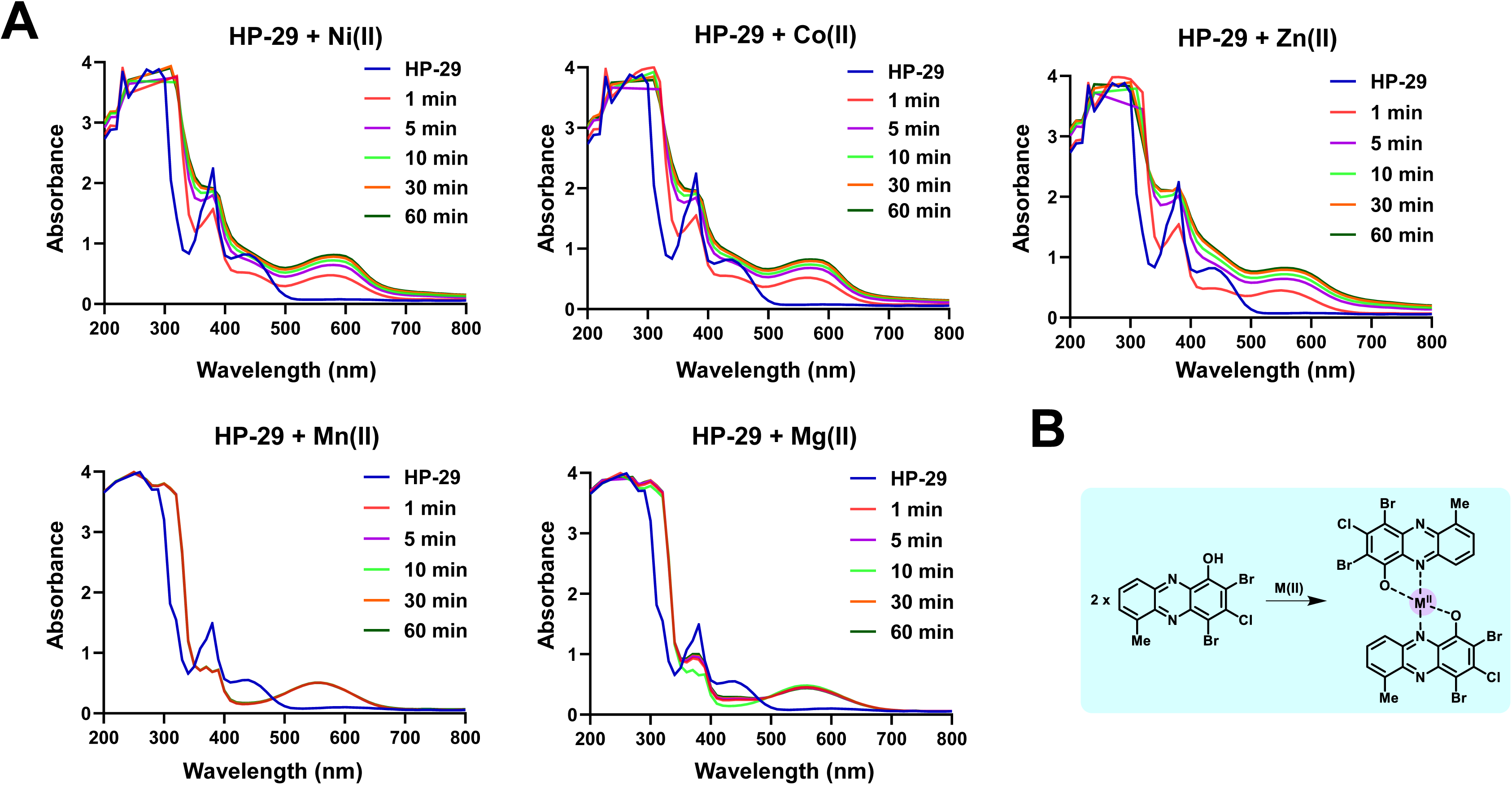
HP-29 is able to bind to several divalent metal (II) cations. (A) UV-vis spectroscopy of HP-29 binding nickel (II), cobalt (II), zinc (II), manganese (II),and magnesium (II). (B) There is a 2:1 HP-29:metal (II) cation complex that forms from direct metal binding of HP-29 in UV-vis experiments.

### HP-29 causes a disruption of metal homeostasis in *S. mutans* that is exacerbated by the addition of zinc

The heightened potency of HP-29 in the presence of zinc, combined with the confirmation that HP-29 is capable of binding to a range of divalent metal cations, introduced the possibility that the inhibitory effect of HPs may be associated with a more extensive disruption of metal homeostasis. To assess the broader impact of HP-29 on intracellular metal content, mid-exponential phase cultures of *S. mutans* UA159 were grown in BHI and exposed to sub-inhibitory concentrations of (i) 0.025 μM HP-29, (ii) 0.5 mM ZnSO_4_, or (iii) 0.025 μM HP-29 and 0.5 mM ZnSO_4_ for 90 minutes, with each treatment group individually compared to a control condition kept in BHI. As expected, treatment with zinc alone did not significantly affect intracellular zinc levels (**Fig. 6A**) as *S. mutans* can maintain zinc homeostasis in high concentrations of zinc due to the activity of the ZccE exporter (21). As a result, treatment with zinc did not impact the intracellular levels of the other metals (iron, manganese, cobalt, nickel and magnesium) tested (**Fig. 6B-F**). Treatment with HP-29 alone resulted in a significant decrease in intracellular iron content that dropped ∼ 40% when compared to the untreated control (**Fig. 6B**). Though not significant, the dual HP-29 and zinc treatment revealed an additional 10% decrease in intracellular iron as compared to the treatment with only HP-29. These results provide the first direct evidence that HP-29 disrupts intracellular iron homeostasis in bacteria. While HP-29 did not affect the intracellular levels of zinc, manganese, nickel or magnesium, exposure to HP-29 alone or combined with zinc caused a significant 2-fold increase in intracellular cobalt (**Fig. 6D**). In addition to iron, dual-treatment with zinc and HP-29 revealed other important changes in intracellular metal content. Most strikingly, intracellular zinc soared in the dual-treatment condition, to levels that were more than 3-fold greater than those seen in either of the single compound treatments (**Fig. 6A**). Furthermore, dual treatment decreased intracellular manganese by ∼ 50% and magnesium by ∼ 30% with no significant changes in cobalt or nickel content observed when compared to HP-29 alone (**Fig. 6C-E**). Notably, the dual treatment significantly disrupted the zinc-to-manganese ratio, reversing the typical balance that favors manganese by approximately 50% to instead favor zinc in an 8:1 ratio (**Fig. 6G**). This shift reflects the well-documented phenomenon in which zinc-induced manganese deficiency broadly impacts bacterial physiology (21, 41, 42).

**Fig. 6.**
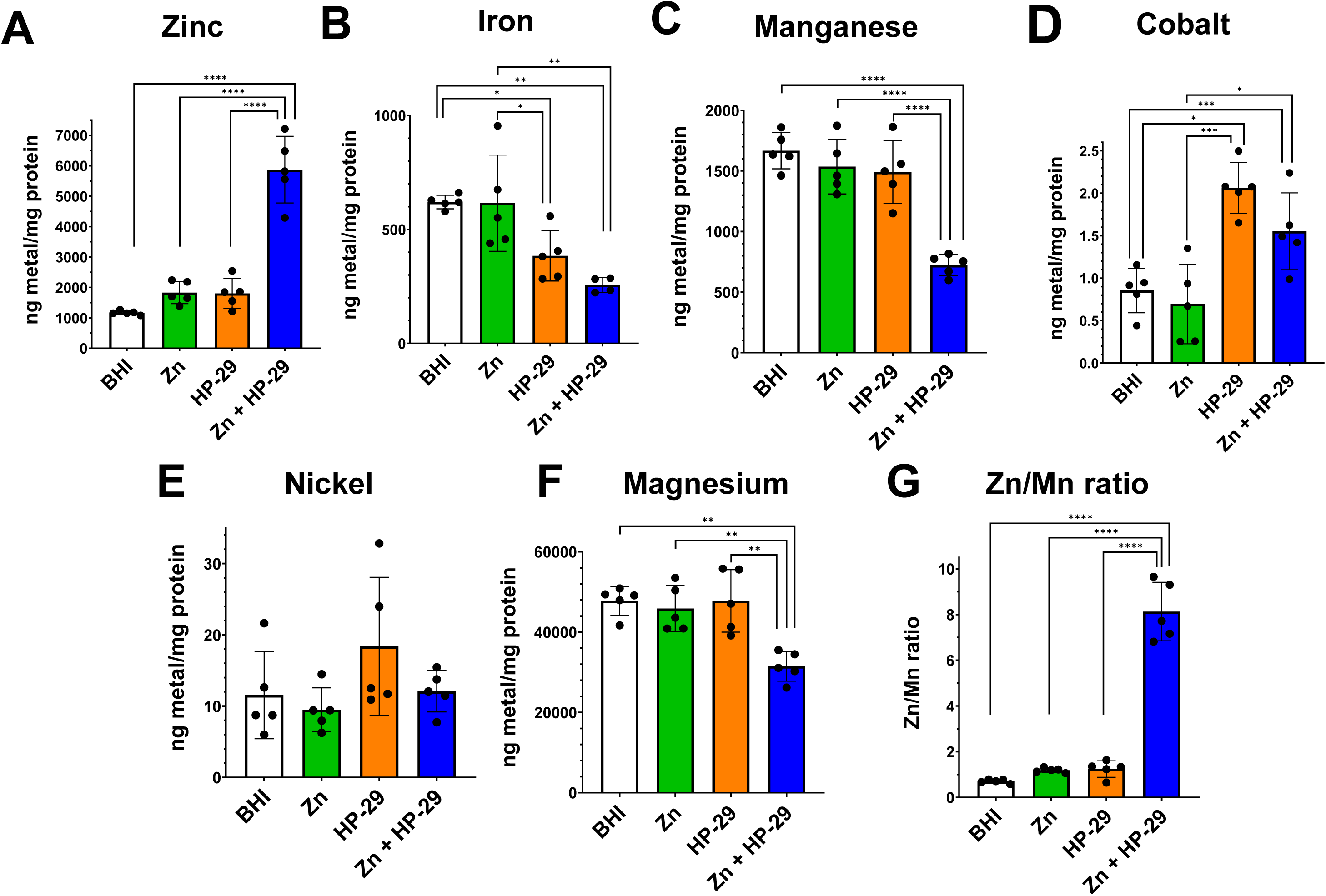
ICP-MS quantification of intracellular biometals reveals that exposure to HP-29 disrupts metal homeostasis in *S. mutans*. Mid-exponential phase cultures (OD_600_ = 0.4) of *S. mutans* UA159 were exposed to BHI alone or supplemented with 0.5 mM ZnSO_4_, 0.025 μM HP-29, or both for 90 minutes. After washing in PBS, the ICP-MS analysis was performed on the harvested cells to determine the intracellular content of biometals: (A) zinc, (B) iron, (C) manganese, (D) nickel, (E) cobalt, or (F) magnesium. (G) shows the intracellular zinc:manganese ratio in each condition. Data represent averages and standard deviations of 5 independent experiments.

### Transcriptome analysis following exposure to HP-29 clearly indicates trace metal stress

To obtain clues on the antimicrobial effects of HP-29 and better understand the synergistic association of HP-29 with zinc, *S. mutans* UA159 cultures were grown to mid-log phase and treated for 30 minutes with 0.025 μM HP-29 or 0.025 μM HP-29 and 0.5 mM ZnSO_4_ and subjected to RNA sequencing (RNA-seq) analysis. When compared to untreated cells kept in BHI, only 12 genes were significantly altered in expression after applying a 2-fold linear cutoff (1-fold log_2_) (**Fig. 7A**, **Table 2**; *p* < 0.05). Yet, the identity of these 12 (10 up and 2 downregulated) genes reveals a compelling story of disruption of metal homeostasis. Among the upregulated genes was the entire *sloABC* operon and their cognate regulator *sloR* (4.7 to 6.3-fold linear; 2.3 to 2.6-fold log_2_), and *mntH* (2.3-fold linear; 1.2-fold log_2_). The *sloABC* operon encodes a highly conserved ABC transporter that mediates iron and manganese uptake, while *mntH* encodes an Nramp-type transporter that mediates manganese uptake (25, 26). In addition, the zinc exporter *zccE* was upregulated by 6.8-fold linear (2.8-fold log_2_) revealing that HP-29 triggers a high zinc stress response even when cells are grown in a low-zinc medium such as BHI (25). The most highly upregulated genes were *smu.236c*, *smu.237c*, and *smu.238c* (from 13 to 46-fold linear; 3.6 to 5.5-fold log_2_), an uncharacterized 3-gene operon comprised of a TetR-type regulator (*smu.236c*), as well as permease (*smu.237c*) and ATP-binding (*smu.238c*) proteins of an ABC transporter (43). Among the only two downregulated genes were *dpr* (0.5-fold linear; –1.0-fold log_2_) and *smu.635* (0.4-fold linear; –1.2-fold log_2_). The PerR-regulated *dpr* gene encodes a ferritin-like protein that acts as an Fe^2+^ sink, protecting cells from oxidative stress caused by Fenton chemistry (20, 44). The downregulation of *dpr* further suggests that HP-29 induces iron starvation. Additionally, *smu.635* encodes an uncharacterized transmembrane protein; although its function is unknown, it is also a member of the PerR regulon (34). Homology predictions suggest that *smu.635* may be involved in manganese transport and homeostasis (34, 43).

**Fig. 7.**
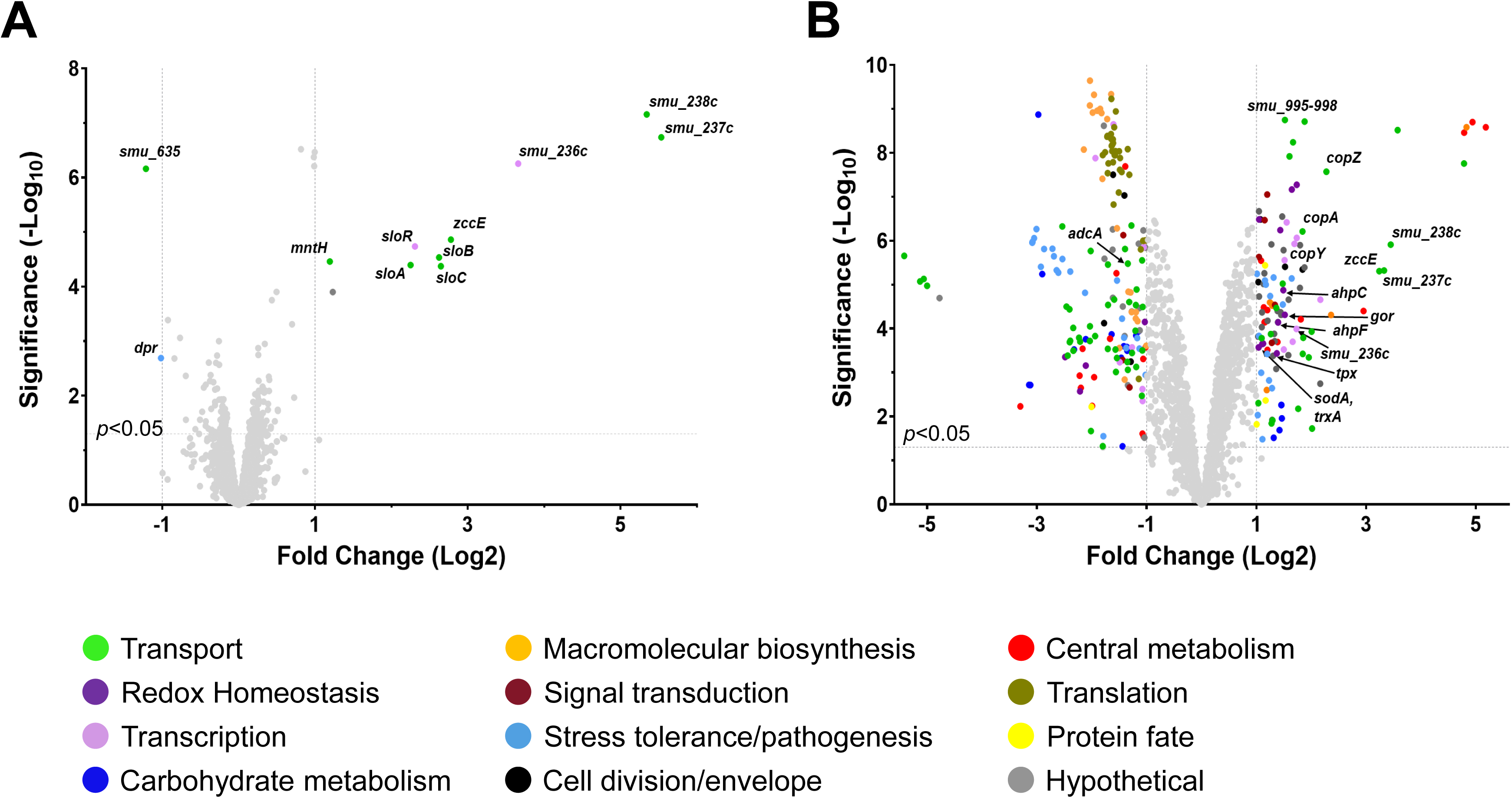
Volcano plots of genes differentially expressed in *S. mutans* UA159 following a 30 minute exposure to. (A) 0.025 µM HP-29 or (B) 0.025 µM HP-29 and 0.5 mM ZnSO_4_ as compared to a BHI control condition. The x-axes indicate the log_2_ fold change in expression while the y-axes indicate the significance. Selected genes of interest are noted. Colors are used to indicate the predicted function of the genes as listed. Genes that do not meet the threshold for significance (1 fold change, *p*<0.05), are shown in light gray.

**Table 2.** *S. mutans* differentially expressed genes when grown in BHI with 0.025µM HP-29 as compared to BHI medium.

| Locus | Gene | Description | Fold change, Log2 | p-value |
| --- | --- | --- | --- | --- |
| SMU_182 | *sloA* | metal ABC transporter ATP-binding protein | 2.2507 | 4.04E-05 |
| SMU_183 | *sloB* | metal ABC transporter permease | 2.6239 | 2.93E-05 |
| SMU_184 | *sloC* | metal ABC transporter substrate-binding protein | 2.6482 | 4.23E-05 |
| SMU_186 | *sloR* | metal-dependent transcriptional regulator SloR | 2.3081 | 1.84E-05 |
| SMU_236c |  | TetR/AcrR family transcriptional regulator | 3.6597 | 5.57E-07 |
| SMU_237c |  | ABC transporter permease | 5.5329 | 1.84E-07 |
| SMU_238c |  | ABC transporter ATP-binding protein | 5.342 | 6.99E-08 |
| SMU_540 | *dpr* | DNA starvation/stationary phase protection protein | -1.017 | 2.05E-03 |
| SMU_635 |  | VIT family protein | -1.2144 | 6.92E-07 |
| SMU_768c |  | hypothetical protein | 1.2317 | 1.26E-04 |
| SMU_770c | *mntH* | Nramp family divalent metal transporter | 1.194 | 3.47E-05 |
| SMU_2057c | *zccE* | heavy metal translocating P-type ATPase | 2.7787 | 1.38E-05 |

Exposure to the dual HP-29/zinc combination resulted in a much greater number of genes differentially expressed as compared to the untreated condition, with 119 genes upregulated and 182 genes downregulated (**Fig. 7B**, **Table S1**, 1-fold cutoff (log_2_), *p* < 0.05). Not surprisingly, dual treatment increased *zccE* expression by 9.4-fold linear (3.2-fold log_2_) while transcription of *adcA*, coding for the substrate binding protein of the zinc uptake system AdcABC (22), was downregulated by 2.4-fold linear (–1.3-fold log_2_). The genes comprising the *smu.995-998* operon, which we have previously described to be involved in iron uptake (20), also showed increased expression (2.9 to 3.7-fold linear; 1.5 to 1.9-fold log_2_) following dual treatment. However, the iron-manganese transporter *sloABC* and manganese transporter *mntH*, induced after HP-29 treatment, were not found to be significantly upregulated in the dual treatment. Also significantly upregulated were the genes of the *copYAZ* operon (3.2 to 4.8-fold linear; 1.7 to 2.3-fold log_2_) that mediate copper export. Finally, the genes of the *smu.236c-238c* operon were also strongly upregulated (3.3 to 10.9-fold linear; 1.7 to 3.4-fold log_2_) following dual treatment. Of note, the *smu.236c-238c* operon was not impacted in our previous study of the *S. mutans* transcriptome following exposure to zinc (21), indicating that this induction is driven by HP-29. In total, 66 transport-associated genes were differentially expressed following the dual treatment exposure. As previously observed under zinc stress alone (21), several genes associated with oxidative stress tolerance were also upregulated upon dual treatment, including *ahpCF*, *gor*, *gst*, *tpx*, *gloA*, *sodA*, and *trxA*. For comparison purposes, Table S1 containing the complete list of genes that experienced significant changes in expression following treatment with both HP-29 and zinc, indicates which of these genes were also identified in our earlier study of the impact of zinc on the *S. mutans* transcriptome. Overall, the dual treatment transcriptome analysis reenforces the notion that HP-29 causes broad disruption of trace metal homeostasis, and that the effect is exacerbated in the presence of elevated, but not inherently inhibitory, zinc concentrations.

To validate the results obtained in the RNA-seq analysis, qRT-PCR was utilized to examine expression of selected metal transport genes after HP-29 (0.025 μM HP-29) and HP-29/zinc (0.025 μM HP-29 + 0.5 mM ZnSO_4_) exposure but this time a zinc-only (0.5 mM ZnSO_4_) treatment was included, which allowed comparisons of the impact of individual and dual treatments directly to each other (**Fig. 8**). As anticipated, expression of *zccE* was significantly induced (∼10-fold) in all three treatment conditions, as compared to the untreated controls, while only the combination of HP-29 and zinc resulted in a significant increase in expression of the copper exporter *copA*. The qRT-PCR profile of the dual transporter *sloC* and manganese transporter *mntH* mirrored the RNA-seq results as HP-29 treatment alone led to significant upregulation of both transporters while dual treatment did not. Finally, the elevated expression of *smu.237c* (ABC transport permease) following exposure to HP-29 or both HP-29 and zinc, but not zinc alone, was confirmed by qRT-PCR.

**Fig. 8.**
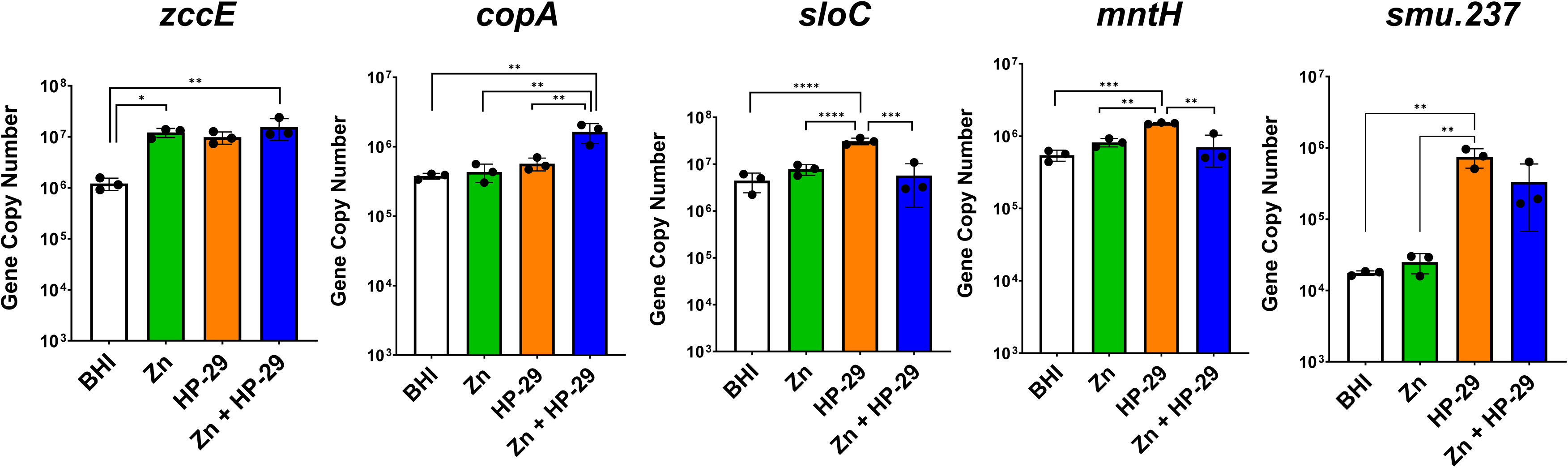
qRT-PCR analysis of genes associated with metal homeostasis in *S. mutans* UA159 following a 30 minute exposure to 0.025 µM HP-29, 0.5 mM ZnSO_4_, or both. *zccE* and *copA* encode proteins associated with zinc efflux. *sloC* and *mntH* encode proteins that function in iron (*sloC*) and manganese (*sloC* and *mntH*) import. *smu.237* encodes a putative transporter of unknown function that is highly upregulated following exposure to HP-29. Data represent averages and standard deviations of at least 3 independent experiments. ****, *p* ≤ 0.0001, ***, *p* ≤ 0.001, **, *p* ≤ 0.01, *, *p* ≤ 0.05.

## DISCUSSION

Antimicrobial resistance is a major global concern, as introduction of antibiotics into clinical use is often (and quickly) followed by the emergence of MDR bacterial strains, placing an enormous burden on healthcare worldwide (2, 45). Halogenated phenazines (HP) were initially discovered from efforts inspired by phenazine antibiotics produced by *P. aeruginosa* in biological warfare against *S. aureus* (4, 6, 7, 9). One potent HP biofilm-eradicating agent discovered during initial efforts, HP-14, was shown to cause a rapid iron starvation in MRSA biofilms (11). Further structural modifications of the original HP scaffold led to the identification of HP-29, which demonstrates greater antibacterial and biofilm-killing activities (10).

Earlier studies have demonstrated that HP-29 is an effective antimicrobial against several bacterial species including *S. aureus*, *S. epidermidis*, *E. faecalis*, and *S. pneumoniae* (10), but neither HP-29 nor other HP analogs have been tested against the oral pathobiont *S. mutans*. Given our long-standing interest in the mechanisms of metal ion homeostasis in *S. mutans* and its link to virulence, this study investigated the impact of HP-29 on iron homeostasis in this major dental and occasional systemic pathogen. As we demonstrate the potential usefulness of HP-29 as an antimicrobial to combat *S. mutans* infections, we discovered that divalent metals including cobalt, iron, manganese and nickel rescued *S. mutans* from HP-29– induced growth inhibition to varying degrees. Unexpectedly, sub-inhibitory concentrations of zinc significantly enhanced the antimicrobial activity of HP-29. This HP-29 plus zinc synergistic effect was also observed in at least two Gram-positive pathogens (*E. faecalis* and *S. aureus*) commonly associated with MDR infections, suggesting that combined HP-29 and zinc treatment may represent a novel therapeutic strategy for broad-spectrum bacterial control.

Through intracellular metal quantification and transcriptome analyses, we demonstrate that HP-29 broadly disrupts metal homeostasis, extending beyond the previously reported role in inducing iron starvation (10, 11). For the first time, we show that HP-29 treatment significantly reduces intracellular iron levels in bacteria. Additionally, our ICP-MS analysis reveals that HP-29 increases intracellular cobalt concentrations, while exerting modest effects on zinc and manganese pools, elevating zinc levels and reducing manganese levels. As ZccE has been shown to mediate tolerance to both zinc and cobalt (21), the increased transcription of *zccE* following HP-29 treatment could be a response to the influx of both metals. Perhaps the most compelling result was a striking 5-fold increase in intracellular zinc in cells treated with both HP-29 and zinc, as compared to its untreated counterpart. This dual HP-29/zinc treatment also resulted in intracellular zinc quantities 3-fold greater than after treatment with either compound individually. A well-established consequence of high intracellular zinc stress is depletion of manganese (21, 41, 42, 46), and indeed manganese levels dropped by approximately 50% in the dual-treated cells compared to single treatment or untreated groups. As a result, these cells experienced a severe disruption in intracellular manganese:zinc balance, which is expected to negatively impact multiple cellular functions, including oxidative stress tolerance (21, 41, 42). In *S. mutans* and related bacteria, manganese contributes to oxidative stress survival by serving as cofactor of the manganese-dependent superoxide dismutase (SOD) enzyme (47, 48) and, likely, by replacing iron as an enzymatic co-factor thereby protecting iron-binding proteins from Fenton chemistry-induced damage. These findings provide a direct explanation for why manganese supplementation alleviates the antimicrobial effects of HP-29. Furthermore, several genes associated with oxidative stress management were upregulated in dual-treated cells, supporting previous findings that zinc mismetallation is a key trigger of oxidative stress (21).

Although not a transition metal, magnesium levels were also measured following treatment. As the most abundant cation in living cells, magnesium plays a critical role in numerous cellular functions, including membrane stabilization, oligonucleotide folding, enzymatic cofactor activity, and participation in stress responses and virulence (49, 50). While individual treatments with HP-29 or zinc had no significant effect on intracellular magnesium levels, dual treatment resulted in a 30% decrease in magnesium content. This sharp decline is likely to impair multiple essential cellular processes, thereby contributing to the heightened susceptibility to HP-29. For example, studies in the model Gram-positive organism *B. subtilis* have shown that magnesium-depleted cells are unable to maintain normal protein translation and enter a state of stasis until magnesium homeostasis is restored (51).

Iron, manganese, and zinc are particularly important during infection, as the host actively limits pathogen access to these essential metals by producing metal-chelating proteins. In response, bacteria have evolved sophisticated mechanisms to scavenge these metals from host tissues (36, 52–54). Our transcriptome analysis strongly supported the conclusion that HP-29 broadly disrupts trace metal homeostasis. Focusing first on HP-29 exposure alone, we observed differential expression of a dozen genes (using a 2-fold linear change cutoff), with the majority clearly associated with metal transport and homeostasis. Notably, genes encoding manganese import systems, *sloABC* and *mntH*, were upregulated, while *zccE* was downregulated. When the cutoff was relaxed to include genes with a 1.5-fold linear change, we also detected upregulation (∼1.8-fold) of the *smu.995–998* operon, which is implicated in iron transport as is the dual transporter *sloABC* (20). Collectively, this transcriptional response suggests that *S. mutans* activates multiple pathways to restore iron, manganese, and zinc homeostasis following HP-29 treatment. Despite using a much lower concentration of zinc (0.5 mM ZnSO₄ vs. 4 mM in our previous transcriptome study), the dual-treatment transcriptome still exhibits key features of the zinc stress response. These include upregulation of metal exporters *zccE* and *copA*, oxidative stress genes, and two major operons involved in lactose uptake and utilization. Notably, *smu.236c-238c*, an uncharacterized TetR regulator and 2-gene ABC transporter lacking a cognate substrate-binding protein-encoding gene, was the most highly upregulated transcriptional unit in HP-29-treated cells and among the most upregulated in dual-treated cells. ABC transporters without substrate-binding proteins are not uncommon and have been linked to molecule export more often than import. Further investigation into the potential role of *smu.236c-238c* in metal transport is warranted.

In summary, here we expand the breadth of Gram-positive pathogens that are susceptible to HPs, specifically HP-29, to include three species of oral streptococci. While confirming that HP-29 induces iron starvation, we further demonstrate that it disrupts broader aspects of metal homeostasis, as evidenced by altered intracellular levels of cobalt and manganese. Additionally, unlike iron and manganese, which can rescue HP-29 sensitivity, zinc acts synergistically with HP-29. This suggests that a combined HP-zinc therapy may offer a promising strategy for treating infections caused by Gram-positive pathogens.

## MATERIALS AND METHODS

### Bacterial strains and growth conditions

The bacterial strains used in this study are listed in Table 3. Oral streptococci were routinely grown in BHI broth at 37°C in a 5% CO_2_ atmosphere. For physiologic analyses, bacterial inocula were prepared from overnight cultures, then sub-cultured 1:20 into fresh media and grown to mid-logarithmic phase (OD_600_ of 0.4) and then diluted 1:25 into the indicated medium (BHI +/-HP-29 and +/-divalent metals) in a microtiter plate. Growth was monitored using the BioScreenC growth reader (Growth Curves USA) at 37°C with each well overlayed with sterile mineral oil to minimize oxidative stress once the plate is shaken prior to OD_600_ readings. For RNA-Seq analysis, replicate cultures of *S. mutans* UA159 were grown as described above to OD_600_ of 0.4 and separated into four aliquots: (i) BHI control, (ii) 0.5 mM ZnSO_4_, (iii) 0.025 μM HP-29, and (iv) 0.5 mM ZnSO_4_ + 0.025μM HP-29. These samples were incubated for an additional 30 minutes, then harvested by centrifugation, and then bacterial pellets resuspended in 1 mL RNA Protect Bacterial Reagent (Qiagen). Following another centrifugation cycle, the supernatants were discarded and the pellets stored at –80°C until use. For ICP-MS analysis, the cultures were grown as described above for RNA-Seq analysis but incubated for 90 minutes upon addition of ZnSO_4_ and/or HP-29. The bacterial cells were harvested by centrifugation, then washed twice in PBS. After a final round of centrifugation, the harvested cell pellets were stored at –20°C until use. Similar methods were used for growth of *E. faecalis* and *S. aureus*, but these strains were incubated at 37°C in an aerobic environment.

**Table 3.** Bacterial strains used in this study.

| Strain | Relevant characteristics | Source or reference |
| --- | --- | --- |
| *Streptococcus mutans* UA159 | Wild-type | ATCC |
| *Streptococcus mutans* $\Delta zccE$ | *zccE*::Spec | (21) |
| *Streptococcus gordonii* DL-1 | Wild-type | ATCC |
| *Streptococcus sanguinis* SK36 | Wild-type | ATCC |
| *Enterococcus faecalis* OG1RF | Wild-type | ATCC |
| *Staphylococcus aureus* RN4220 | Wild-type | BEI Resources |

### MIC assays

The minimum inhibitory concentration (MIC) of HP-29 was determined by a broth microdilution method using two-fold serial dilutions of HP-29, of ≥ 95% purity (21). Briefly, mid-logarithmic phase cultures (OD_600_ of 0.4, 1.5 x 10^9^ CFU/mL) were diluted 1:25 in the indicated medium. 96-well plates were incubated at 37°C in the appropriate atmosphere for 24 hours and the concentration of HP-29 at which the absorbance values were 15% or less of the control condition was determined to be the MIC. Assays were performed with a minimum of three biological replicates.

### Metal binding to HP-29 (UV-vis)

A mixture of DMSO (970 μL) and HP-29 (30 μL of a 10 mM DMSO stock) was added to a 1.5 mL cuvette. In a separate cuvette, DMSO (955 μL), HP-29 (30 μL of a 10 mM DMSO stock), and metal (II) cation (15 μL of a 10 mM water solution) were added and thoroughly mixed. Then, spectral scanning was performed from 200 to 800 nm in 10 nm increments at various time points (1, 5, 10, 30, & 60 min). All divalent metal salt solutions were freshly made and added to the cuvette immediately. The elevated absorbance between 500 – 700 nm of HP sample after the addition of divalent metal indicates rapid binding between HP-29 and metal. Metal salts used during these studies: ZnSO_4_, NiSO_4_, CoSO_4_, MgSO_4_, and MnSO_4_·H_2_O.

### ICP-MS analysis

The intracellular metal content of *S. mutans* UA159 was determined via inductively coupled plasma mass spectrometry (ICP-MS) performed at the University of Florida Analytical Toxicology Core Laboratory (ATCL). The cell pellets were resuspended in HNO_3_ and incubated at 100°C for 30 minutes. Metal concentrations were determined using an Agilent 7900 ICP MS equipped with in-line internal standard addition. Data acquisition was accomplished in helium gas mode. Calibration ranges for all metals were 0-10,000 ng/mL, and all linear regression r^2^ values were 1.0000. Metal concentrations were normalized to total protein content determined by the bicinchoninic acid (BCA) assay (Pierce). The data shown were collected from 5 biological replicates. Analysis of variance (ANOVA) was performed to verify significance of the results.

### RNA sequencing analysis

Total RNA was isolated from homogenized *S. mutans* cell lysates by acid-phenol:chloroform extractions as previously described (34). Briefly, nucleic acid obtained after homogenization was digested with Ambion DNaseI (Thermo Fisher), then purified using the RNeasy kit (Qiagen), which included an on-column DNase digestion according to manufacturer’s instructions. Sample quality and quantity were assessed on an Agilent 2200 Tape Station at the University of Florida Interdisciplinary Center for Biotechnology Research (ICBR). For RNA-Seq analysis, purified samples were sent to SeqCenter (Pittsburgh, PA) for DNase treatment. rRNA depletion was performed with the Ribo-Zero Plus Microbiome kit (Illumina) to generate RNA libraries. Following cDNA synthesis, libraries were subjected to RNA deep sequencing using the Illumina NovaSeq platform. Read quantification was performed using Subread’s featureCounts2 functionality. Quality control and adapter trimmer were performed with bcl-convert. Read mapping was performed with HISAT2 using the *S. mutans* UA159 genome (GenBank accession number NC_004350.2). Read counts loaded into R were normalized using edgeR’s Trimmed Mean of M values (TMM) algorithm. Subsequent values were then converted to counts per million (CPM). Differential expression analysis was performed using edgeR’s glmQLFTest.

Targeted gene expression analysis was performed by quantification of mRNA by quantitative reverse transcriptase real-time PCR (qRT-PCR), performed according to an established protocol (24) using gene-specific primers listed in Table 4. Analysis of variance (ANOVA) was performed to verify significance of the results.

**Table 4.**
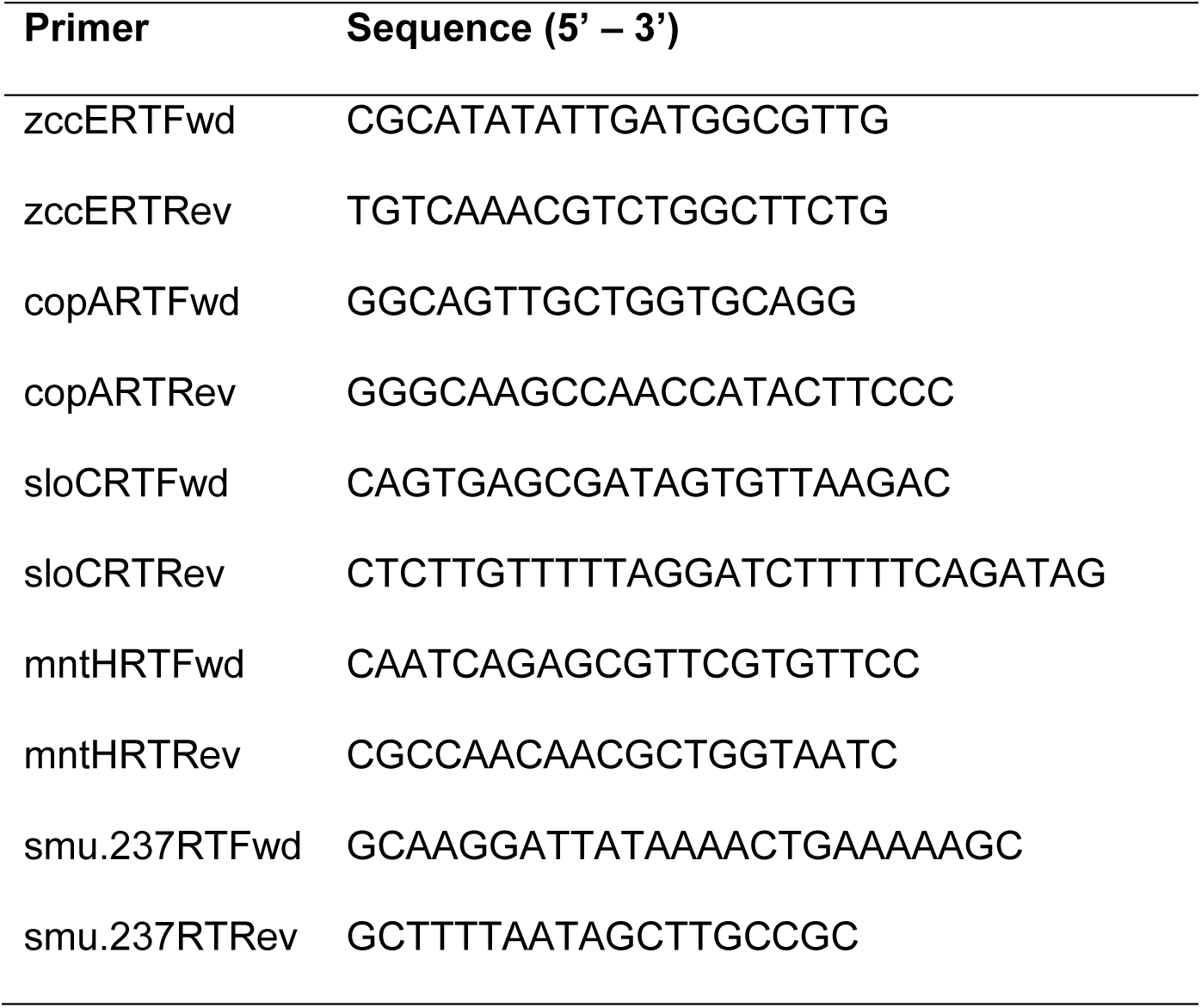
qRT-PCR primers used in this study.

## Data availability

Gene expression data have been deposited in the NCBI Gene Expression Omnibus (GEO) database (https://www.ncbi.nlm.nih.gov/geo/) under GEO Series accession number GSE296169.

## ACKNOWLEDGEMENTS

This study was supported by NIH-NIDCR R01 DE032555 to J.A.L. and NIH-GMS R35 GM153272 to R.W.H.

