## Supplemental Table 1 for "Zinc-Enhanced Activity of an Antimicrobial Halogenated Phenazine Against *Streptococcus mutans* and Other Gram-positive Bacteria"

**Table S1** *S. mutans* differentially expressed genes when grown in BHI with 0.025 µM HP-29 and 0.5 mM Zinc as compared to BHI medium.

| Locus | Gene | Description | Fold change, Log2 | *p*-value | Observed in zinc exposure study? (1) |
|---|---|---|---|---|---|
| Upregulated genes | | | | | |
| SMU_05 | | DUF951 domain-containing protein | 1.4703 | 2.83E-07 | Yes |
| SMU_143c | def | peptide deformylase | 1.1577 | 3.65E-06 | |
| SMU_144c | | cyclic nucleotide-binding domain-containing protein | 1.5123 | 2.79E-06 | |
| SMU_145 | | MFS transporter | 1.2601 | 1.33E-04 | |
| SMU_236c | | TetR/AcrR family transcriptional regulator | 1.7302 | 1.02E-04 | |
| SMU_237c | | ABC transporter permease | 3.3266 | 4.76E-06 | |
| SMU_238c | | ABC transporter ATP-binding protein | 3.4473 | 1.23E-06 | |
| SMU_239c | | VanZ family protein | 1.4490 | 4.87E-05 | Yes |
| SMU_363 | | MerR family transcriptional regulator | 2.1643 | 2.20E-05 | Yes |
| SMU_364 | glnA | type I glutamate--ammonia ligase | 2.3602 | 4.90E-05 | Yes |
| SMU_365 | gltB | glutamate synthase large subunit | 1.1874 | 2.51E-03 | Yes |
| SMU_383c | | SDR family oxidoreductase | 1.1039 | 2.25E-04 | |
| SMU_424 | copY | CopY/TcrY family copper transport repressor | 1.6905 | 1.18E-06 | Yes |
| SMU_426 | copA | heavy metal translocating P-type ATPase | 1.8385 | 6.15E-07 | Yes |
| SMU_427 | copZ | copper chaperone CopZ | 2.2749 | 2.70E-08 | Yes |
| SMU_503c | | hypothetical protein | 1.7932 | 1.19E-05 | Yes |
| SMU_561c | | NUDIX hydrolase | 1.3835 | 2.03E-04 | Yes |
| SMU_602 | | bile acid:sodium symporter family protein | 1.2935 | 1.22E-02 | Yes |
| SMU_609 | | SH3 domain-containing protein | 1.5204 | 3.92E-06 | Yes |
| SMU_610 | spaP | cell surface antigen I/II | 1.8388 | 4.52E-06 | |
| SMU_629 | sodA | superoxide dismutase SodA | 1.0397 | 2.69E-04 | Yes |
| SMU_647 | | O-methyltransferase | 1.1339 | 3.31E-05 | Yes |
| SMU_670 | acnA | aconitate hydratase AcnA | 5.1800 | 2.62E-09 | Yes |
| SMU_671 | | citrate synthase | 4.9365 | 2.00E-09 | Yes |

| | | | | | |
|---|---|---|---|---|---|
| SMU_672 | icd | NADP-dependent isocitrate dehydrogenase | 4.7856 | 3.47E-09 | Yes |
| SMU_673 | | putative ABC transporter permease | 3.5720 | 3.06E-09 | Yes |
| SMU_724 | | glycerophosphodiester phosphodiesterase | 1.0352 | 8.76E-06 | Yes |
| SMU_727 | | MerR family transcriptional regulator | 1.5508 | 3.82E-07 | Yes |
| SMU_728 | | NADP-dependent oxidoreductase | 1.6433 | 6.86E-08 | Yes |
| SMU_730 | | AbrB/MazE/SpoVT family DNA-binding domain-containing protein | 1.5810 | 4.05E-04 | |
| SMU_731 | | ABC transporter ATP-binding protein | 1.0354 | 4.96E-03 | |
| SMU_764 | ahpC | alkyl hydroperoxide reductase subunit C | 1.4888 | 1.33E-05 | Yes |
| SMU_765 | ahpF | alkyl hydroperoxide reductase subunit F | 1.3951 | 7.23E-05 | Yes |
| SMU_805c | | amino acid ABC transporter ATP-binding protein | 1.3623 | 3.40E-05 | Yes |
| SMU_806c | | ABC transporter substrate-binding protein/permease | 1.4792 | 9.54E-06 | Yes |
| SMU_838 | gorA | glutathione-disulfide reductase | 1.5155 | 4.87E-05 | Yes |
| SMU_839 | | folylpolyglutamate synthase/dihydrofolate synthase family protein | 1.0469 | 2.14E-07 | Yes |
| SMU_911c | | DUF308 domain-containing protein | 1.7936 | 1.27E-06 | Yes |
| SMU_913 | gdhA | NADP-specific glutamate dehydrogenase | 1.2891 | 2.84E-05 | Yes |
| SMU_924 | tpx | thiol peroxidase | 1.3705 | 3.66E-04 | Yes |
| SMU_925 | | hypothetical protein | 1.4413 | 4.28E-05 | Yes |
| SMU_984 | | CHAP domain-containing protein | 1.8744 | 4.06E-06 | Yes |
| SMU_991 | | ATP cone domain-containing protein | 1.3025 | 4.18E-04 | Yes |
| SMU_995 | | ABC transporter permease; iron chelate uptake ABC transporter family permease subunit | 1.5195 | 1.80E-09 | Yes |
| SMU_996 | | iron chelate uptake ABC transporter family permease subunit | 1.6021 | 1.21E-08 | Yes |
| SMU_997 | | ABC transporter ATP-binding protein | 1.6657 | 5.75E-09 | Yes |
| SMU_998 | | siderophore ABC transporter substrate-binding protein | 1.8776 | 1.95E-09 | Yes |
| SMU_999 | | hypothetical protein | 1.3852 | 3.98E-05 | Yes |
| SMU_1009 | | sensor histidine kinase | 1.0455 | 2.36E-06 | |
| SMU_1036 | | ABC transporter permease | 1.0943 | 1.65E-04 | |
| SMU_1037c | | sensor histidine kinase KdpD | 1.1970 | 8.89E-08 | |
| SMU_1038c | | response regulator transcription factor | 1.1486 | 3.40E-07 | Yes |

| | | | | | |
|---|---|---|---|---|---|
| SMU_1039c | | glycosyltransferase family 8 protein | 1.0738 | 3.29E-07 | Yes |
| SMU_1040c | | SDR family oxidoreductase | 1.0486 | 3.29E-07 | Yes |
| SMU_1048 | | CYTH domain-containing protein | 1.1296 | 9.65E-06 | Yes |
| SMU_1071c | | alpha/beta hydrolase | 1.1475 | 7.16E-05 | |
| SMU_1072c | | GNAT family N-acetyltransferase | 2.9509 | 3.98E-05 | |
| SMU_1073 | | formate--tetrahydrofolate ligase | 1.1972 | 3.80E-05 | |
| SMU_1175 | | sodium:alanine symporter family protein | 2.0142 | 1.89E-02 | Yes |
| SMU_1259 | | Eco57I restriction-modification methylase domain-containing protein | 1.0866 | 1.01E-03 | |
| SMU_1287 | | MerR family transcriptional regulator | 1.4986 | 3.00E-04 | Yes |
| SMU_1294 | | flavodoxin | 1.0830 | 2.85E-06 | |
| SMU_1296 | yghU | glutathione-dependent disulfide-bond oxidoreductase | 1.8087 | 6.14E-05 | Yes |
| SMU_1297 | | bifunctional oligoribonuclease/PAP phosphatase NrnA | 1.3242 | 1.32E-04 | Yes |
| SMU_1396 | | glucan-binding protein; GbpC/Spa domain-containing protein | 1.4807 | 2.83E-05 | Yes |
| SMU_1451 | budA | acetolactate decarboxylase | 1.1546 | 7.13E-05 | |
| SMU_1488c | | DUF3884 family protein | 1.2860 | 1.20E-02 | Yes |
| SMU_1489 | | aldose 1-epimerase family protein | 1.4538 | 5.50E-03 | Yes |
| SMU_1490 | lacG | 6-phospho-beta-galactosidase | 1.4547 | 5.47E-03 | Yes |
| SMU_1491 | | lactose-specific PTS transporter subunit EIIC | 1.2750 | 1.44E-02 | Yes |
| SMU_1492 | lacF | PTS lactose transporter subunit IIA | 1.7617 | 6.67E-03 | Yes |
| SMU_1493 | lacD | tagatose-bisphosphate aldolase | 1.4625 | 1.11E-02 | Yes |
| SMU_1494 | lacC | tagatose-6-phosphate kinase | 1.3152 | 3.05E-02 | Yes |
| SMU_1495 | lacB | galactose-6-phosphate isomerase subunit LacB | 1.4203 | 2.04E-02 | Yes |
| SMU_1496 | lacA | galactose-6-phosphate isomerase subunit LacA | 1.0919 | 9.07E-02 | Yes |
| SMU_1498 | lacR | transcriptional regulator LacR | 1.6593 | 2.00E-04 | Yes |
| SMU_1519 | | amino acid ABC transporter ATP-binding protein | 1.8444 | 3.76E-04 | |
| SMU_1520 | | transporter substrate-binding domain-containing protein | 2.0043 | 1.17E-04 | |
| SMU_1521 | | amino acid ABC transporter permease | 1.9538 | 4.53E-04 | |
| SMU_1522 | | amino acid ABC transporter permease | 1.8533 | 1.62E-04 | |
| SMU_1602 | | NAD(P)H-dependent oxidoreductase | 1.4341 | 5.75E-07 | Yes |

| | | | | | |
|---|---|---|---|---|---|
| SMU_1603 | gloA | lactoylglutathione lyase; VOC family protein | 1.7343 | 5.35E-08 | Yes |
| SMU_1649 | | exodeoxyribonuclease III | 1.0461 | 1.47E-04 | |
| SMU_1657c | | P-II family nitrogen regulator | 4.8278 | 2.61E-09 | Yes |
| SMU_1658 | | ammonium transporter | 4.7830 | 1.75E-08 | Yes |
| SMU_1670c | | YlbG family protein | 1.0306 | 1.51E-04 | Yes |
| SMU_1671c | | YlbF family regulator | 1.0970 | 9.29E-05 | Yes |
| SMU_1682c | | type 1 glutamine amidotransferase family protein | 1.2879 | 3.00E-05 | Yes |
| SMU_1700c | cidB | antiholin-like protein; LrgB family protein | 1.3588 | 8.31E-04 | |
| SMU_1704 | | PadR family transcriptional regulator | 1.7332 | 8.63E-07 | Yes |
| SMU_1705 | | DUF1700 domain-containing protein | 1.5861 | 2.20E-05 | Yes |
| SMU_1706 | | DUF4097 family beta strand repeat-containing protein | 1.4942 | 1.65E-06 | Yes |
| SMU_1753c | cas2 | CRISPR-associated endonuclease Cas2 | 1.0091 | 5.69E-06 | Yes |
| SMU_1758c | cas4 | CRISPR-associated protein Cas4 | 1.1722 | 1.00E-05 | Yes |
| SMU_1760c | cas7c | type I-C CRISPR-associated protein Cas7/Csd2 | 1.3129 | 6.86E-06 | Yes |
| SMU_1763c | cas5c | type I-C CRISPR-associated protein Cas5c | 1.1557 | 8.06E-06 | Yes |
| SMU_1764c | | CRISPR-associated helicase/endonuclease Cas3 | 1.2491 | 1.83E-05 | Yes |
| SMU_1803c | | DUF4230 domain-containing protein | 1.1008 | 4.28E-05 | Yes |
| SMU_1862 | | hypothetical protein | 1.2022 | 6.65E-05 | |
| SMU_1865 | mutY | A/G-specific adenine glycosylase | 1.6414 | 7.22E-06 | Yes |
| SMU_1867c | | zinc-dependent alcohol dehydrogenase family protein | 1.2015 | 3.07E-04 | Yes |
| SMU_1869 | trxA | thioredoxin | 1.1212 | 2.20E-04 | Yes |
| SMU_1883 | | DUF956 family protein | 1.0473 | 1.86E-05 | Yes |
| SMU_1916 | | sensor histidine kinase; GHKL domain-containing protein | 1.3364 | 2.90E-05 | |
| SMU_1917 | | response regulator transcription factor | 1.2788 | 2.10E-04 | |
| SMU_1945 | | polyphosphate polymerase domain-containing protein | 1.2728 | 1.21E-06 | |
| SMU_1946 | | DUF4956 domain-containing protein | 1.1446 | 5.51E-06 | |
| SMU_1954 | groL | chaperonin GroEL | 1.0024 | 1.51E-02 | Yes |
| SMU_1955 | groES | co-chaperone GroES | 1.1675 | 4.34E-03 | Yes |
| SMU_1980c | comGG | competence type IV pilus minor pilin ComGG | 1.2304 | 1.51E-03 | |
| SMU_1981c | comGF | competence type IV pilus minor pilin ComGF | 1.1081 | 3.30E-02 | |

| | | | | | |
|---|---|---|---|---|---|
| SMU_1982c | comGE | competence type IV pilus minor pilin ComGE | 1.2829 | 2.27E-03 | |
| SMU_1985 | comGB | competence type IV pilus assembly protein ComGB; type II secretion system F family protein | 1.1965 | 3.77E-04 | |
| SMU_1987 | comGA; tadA | competence type IV pilus ATPase ComGA; Flp pilus assembly complex ATPase component TadA | 1.0231 | 9.28E-03 | |
| SMU_1988c | | DUF1033 family protein | 1.3324 | 1.94E-04 | Yes |
| SMU_2057c | zccE | heavy metal translocating P-type ATPase | 3.2372 | 4.93E-06 | Yes |
| SMU_2074 | nrdD | anaerobic ribonucleoside-triphosphate reductase | 1.2458 | 2.55E-05 | Yes |
| SMU_2129c | | metal-sulfur cluster assembly factor | 1.3921 | 7.17E-05 | Yes |
| SMU_2133c | | YhgE/Pip domain-containing protein | 2.1617 | 1.80E-03 | |
| Downregulated genes | | | | | |
| SMU_16 | | APC family permease | -1.2789 | 4.51E-07 | |
| SMU_20 | mreC | rod shape-determining protein MreC | -1.4076 | 9.33E-08 | |
| SMU_21 | mreD | rod shape-determining protein MreD | -1.7769 | 7.55E-05 | Yes |
| SMU_22 | pcsB | peptidoglycan hydrolase PcsB | -1.6127 | 3.15E-08 | |
| SMU_78 | fruA | fructan beta-fructosidase | -3.1433 | 1.92E-03 | |
| SMU_79 | | glycoside hydrolase family 32 protein | -3.1201 | 1.92E-03 | |
| SMU_89c | | formate/nitrite transporter family protein | -1.3447 | 8.69E-04 | |
| SMU_100 | | PTS sugar transporter subunit IIB | -1.3521 | 5.75E-04 | Yes |
| SMU_101 | | PTS sugar transporter subunit IIC | -1.5436 | 4.66E-04 | Yes |
| SMU_102 | | PTS system mannose/fructose/sorbose family transporter subunit IID | -1.7091 | 2.90E-04 | Yes |
| SMU_103 | | PTS sugar transporter subunit IIA | -1.5615 | 9.68E-04 | Yes |
| SMU_104 | | alpha-glucosidase | -1.9403 | 1.48E-04 | Yes |
| SMU_105 | | LacI family DNA-binding transcriptional regulator | -1.2667 | 2.68E-04 | |
| SMU_133c | | MFS transporter | -1.1840 | 1.29E-05 | |
| SMU_148 | adhE | bifunctional acetaldehyde-CoA/alcohol dehydrogenase | -3.3022 | 5.89E-03 | |
| SMU_179 | | NADPH-dependent FMN reductase | -2.1115 | 6.96E-04 | |
| SMU_180 | | flavocytochrome c | -2.2152 | 2.68E-03 | Yes |
| SMU_263 | | APC family permease | -2.0155 | 2.15E-02 | Yes |
| SMU_264 | aguA | agmatine deiminase | -1.7908 | 2.82E-02 | |
| SMU_265 | arcC | carbamate kinase | -1.7957 | 5.03E-02 | |

| | | | | | |
|---|---|---|---|---|---|
| SMU_292 | | helix-turn-helix transcriptional regulator; AraC family | -1.0539 | 1.41E-06 | Yes |
| SMU_308 | | SDR family oxidoreductase | -1.0345 | 7.07E-05 | |
| SMU_401c | | GNAT family N-acetyltransferase | -1.0201 | 2.67E-04 | |
| SMU_402 | pflB | formate C-acetyltransferase | -2.2237 | 1.18E-03 | Yes |
| SMU_438c | | 2-hydroxyacyl-CoA dehydratase | -1.5518 | 5.53E-06 | |
| SMU_496 | cysK | cysteine synthase A | -1.4018 | 1.46E-03 | |
| SMU_500 | raiA | ribosome hibernation-promoting factor, HPF/YfiA family | -1.1445 | 1.40E-03 | |
| SMU_527 | | dihydrodipicolinate reductase; Gfo/Idh/MocA family oxidoreductase | -1.2734 | 4.20E-05 | |
| SMU_531 | | chorismate mutase | -1.5423 | 5.23E-07 | |
| SMU_532 | trpE | anthranilate synthase component I | -1.3353 | 1.45E-05 | Yes |
| SMU_533 | | aminodeoxychorismate/anthranilate synthase component II | -1.1659 | 6.80E-05 | Yes |
| SMU_534 | trpD | anthranilate phosphoribosyltransferase | -1.2002 | 5.90E-05 | Yes |
| SMU_535 | trpC | indole-3-glycerol phosphate synthase TrpC | -1.0106 | 2.47E-04 | Yes |
| SMU_539c | | A24 family peptidase | -1.2907 | 5.65E-04 | |
| SMU_541 | | YqgQ family protein | -1.4364 | 3.16E-05 | Yes |
| SMU_542 | | ROK family glucokinase | -1.3615 | 2.63E-04 | Yes |
| SMU_543 | | rhodanese-like domain-containing protein | -1.4015 | 1.48E-04 | Yes |
| SMU_574c | lrgB | antiholin-like protein LrgB | -1.9930 | 5.78E-03 | |
| SMU_575c | lrgA | holin-like protein LrgA | -1.9591 | 1.28E-03 | |
| SMU_576 | lytR | two-component system response regulator LytR | -1.3124 | 2.19E-03 | Yes |
| SMU_577 | lytS | two-component system sensor histidine kinase LytS | -1.4386 | 5.22E-04 | Yes |
| SMU_611 | | DEAD/DEAH box helicase | -1.2952 | 1.51E-05 | |
| SMU_616 | | hypothetical protein | -1.6179 | 1.99E-05 | |
| SMU_618 | | hypothetical protein | -1.7692 | 2.57E-06 | |
| SMU_636 | nagB | glucosamine-6-phosphate deaminase NagB | -1.0549 | 2.87E-02 | |
| SMU_675 | ptsP | phosphoenolpyruvate--protein phosphotransferase | -1.2075 | 1.12E-04 | |
| SMU_697 | infC | translation initiation factor IF-3 | -1.0166 | 9.81E-07 | |
| SMU_699 | rplT | 50S ribosomal protein L20 | -1.1085 | 1.58E-06 | |
| SMU_772 | | YSIRK-type signal peptide-containing protein | -1.1262 | 1.10E-04 | |

| | | | | | |
|---|---|---|---|---|---|
| SMU_799c | | thioesterase family protein | -1.3430 | 1.96E-03 | |
| SMU_803c | | ATP-binding cassette domain-containing protein | -1.3958 | 1.54E-06 | |
| SMU_832 | | hypothetical protein | -1.0241 | 1.44E-06 | Yes |
| SMU_865 | rpsP | 30S ribosomal protein S16 | -1.3165 | 3.15E-08 | |
| SMU_866 | | KH domain-containing protein | -1.0604 | 5.83E-07 | |
| SMU_876 | | AraC family transcriptional regulator | -1.0752 | 4.43E-03 | |
| SMU_877 | | alpha-galactosidase | -2.1691 | 2.87E-04 | |
| SMU_878 | msmE | sugar-binding protein MsmE | -2.3942 | 1.90E-04 | Yes |
| SMU_879 | | carbohydrate ABC transporter permease | -2.3377 | 3.20E-04 | Yes |
| SMU_880 | | carbohydrate ABC transporter permease | -2.4045 | 2.03E-04 | Yes |
| SMU_881 | gtfA | sucrose phosphorylase | -2.3388 | 9.65E-05 | Yes |
| SMU_882 | | ABC transporter ATP-binding protein | -2.2131 | 1.99E-04 | Yes |
| SMU_883 | dexB | glucan 1,6-alpha-glucosidase DexB | -2.1114 | 1.75E-04 | Yes |
| SMU_886 | | galactokinase | -1.1933 | 1.44E-04 | |
| SMU_905 | | ABC transporter ATP-binding protein | -1.0735 | 3.22E-05 | Yes |
| SMU_906 | | ABC transporter ATP-binding protein | -1.1979 | 2.92E-05 | Yes |
| SMU_910 | gtfD | glucosyltransferase-S | -2.9764 | 1.35E-09 | Yes |
| SMU_937 | mvaD | diphosphomevalonate decarboxylase | -1.0642 | 4.89E-04 | |
| SMU_957 | rplJ | 50S ribosomal protein L10 | -1.7127 | 2.90E-08 | |
| SMU_960 | rplL | 50S ribosomal protein L7/L12 | -1.6050 | 1.51E-07 | |
| SMU_980 | | beta-glucoside-specific PTS transporter subunit IIABC | -2.4074 | 3.64E-05 | Yes |
| SMU_992 | | hypothetical protein | -1.6219 | 1.61E-06 | Yes |
| SMU_1041 | | ABC transporter ATP-binding protein | -1.0846 | 2.80E-06 | |
| SMU_1042 | | hypothetical protein | -1.1468 | 1.18E-06 | |
| SMU_1062 | | ABC transporter permease/substrate binding protein | -1.7049 | 2.89E-05 | Yes |
| SMU_1063 | | glycine betaine/L-proline ABC transporter ATP-binding protein | -1.5922 | 2.19E-05 | Yes |
| SMU_1070c | | LytTR family DNA-binding domain-containing protein | -1.0413 | 3.04E-02 | |
| SMU_1077 | | phospho-sugar mutase | -2.3240 | 3.01E-04 | Yes |
| SMU_1091 | | putative cross-wall-targeting lipoprotein signal domain-containing protein | -1.7818 | 2.42E-09 | Yes |

| | | | | | |
|---|---|---|---|---|---|
| SMU_1093 | | ABC transporter permease | -1.0788 | 3.14E-04 | |
| SMU_1124 | | pyrimidine-nucleoside phosphorylase | -1.4986 | 2.78E-04 | Yes |
| SMU_1125c | | class I SAM-dependent methyltransferase | -1.6687 | 1.70E-04 | Yes |
| SMU_1185 | | PTS mannitol transporter subunit IICB | -1.8002 | 4.75E-02 | Yes |
| SMU_1284c | | NAD(P)-dependent oxidoreductase | -1.1361 | 3.42E-05 | |
| SMU_1286c | | MFS transporter | -1.3188 | 2.50E-05 | Yes |
| SMU_1302 | adcA | zinc ABC transporter substrate-binding protein AdcA | -1.3449 | 3.34E-06 | Yes |
| SMU_1334 | mubP | mutanobactin A biosynthesis phosphopantetheinyl transferase MubP | -2.6908 | 2.27E-06 | |
| SMU_1335c | mubJ | mutanobactin A biosynthesis reductase MubJ | -2.5377 | 2.62E-06 | Yes |
| SMU_1336 | mubI | mutanobactin A biosynthesis transacylase MubI | -2.7183 | 1.52E-06 | Yes |
| SMU_1337c | mubM | mutanobactin A biosynthesis alpha/beta hydrolase MubM | -2.8711 | 1.54E-06 | |
| SMU_1338c | mubZ | mutanobactin A system MFS transporter MubZ | -3.0089 | 5.45E-07 | |
| SMU_1339 | mubD | mutanobactin A non-ribosomal peptide synthetase MubD | -3.0523 | 8.76E-07 | Yes |
| SMU_1340 | mubC | mutanobactin A non-ribosomal peptide synthetase MubC | -3.0810 | 1.06E-06 | Yes |
| SMU_1341c | mubB | mutanobactin A non-ribosomal peptide synthetase MubB | -3.0851 | 1.11E-06 | Yes |
| SMU_1342 | mubA | mutanobactin A non-ribosomal peptide synthetase MubA | -2.9289 | 3.92E-06 | Yes |
| SMU_1343c | mubH | mutanobactin A polyketide synthase MubH | -2.6412 | 4.70E-06 | Yes |
| SMU_1344c | mubG | mutanobactin A biosynthesis transacylase MubG | -2.6111 | 5.31E-06 | Yes |
| SMU_1345c | mubE | mutanobactin A non-ribosomal peptide synthetase MubE | -2.3929 | 5.05E-06 | Yes |
| SMU_1346 | mubT | mutanobactin A biosynthesis thioesterase MubT | -2.1237 | 1.53E-05 | Yes |
| SMU_1347c | mubY | mutanobactin A system ABC transporter permease subunit MubY | -1.4128 | 1.61E-04 | Yes |
| SMU_1348c | mubX | mutanobactin A system ABC transporter ATP-binding subunit MubX | -1.1738 | 1.64E-04 | Yes |
| SMU_1365c | mubY | mutanobactin A system ABC transporter permease subunit MubY | -1.3729 | 2.76E-04 | Yes |

| | | | | | |
|---|---|---|---|---|---|
| SMU_1366c | mubX | mutanobactin A system ABC transporter ATP-binding subunit MubX | -1.1343 | 2.85E-04 | Yes |
| SMU_1390 | | Pr6Pr family membrane protein | -1.6196 | 5.54E-07 | |
| SMU_1410 | | FAD-dependent oxidoreductase | -2.4865 | 4.45E-04 | |
| SMU_1411 | | MFS transporter | -2.4402 | 4.13E-04 | |
| SMU_1424 | lpdA | dihydrolipoyl dehydrogenase | -1.0765 | 2.47E-02 | |
| SMU_1425 | | ATP-dependent Clp protease ATP-binding subunit; AAA family ATPase | -2.0085 | 6.02E-03 | |
| SMU_1536 | glgA | glycogen synthase GlgA | -1.3154 | 6.09E-02 | Yes |
| SMU_1537 | glgD | glucose-1-phosphate adenylyltransferase subunit GlgD | -1.4409 | 4.81E-02 | Yes |
| SMU_1538 | | glucose-1-phosphate adenylyltransferase | -1.3459 | 5.60E-02 | Yes |
| SMU_1564 | | glycogen/starch/alpha-glucan phosphorylase | -1.3555 | 3.20E-04 | |
| SMU_1565 | malQ | 4-alpha-glucanotransferase | -1.1439 | 1.15E-04 | |
| SMU_1568 | | extracellular solute-binding protein | -1.5707 | 3.00E-04 | |
| SMU_1569 | | carbohydrate ABC transporter permease | -1.0878 | 3.39E-03 | |
| SMU_1570 | | sugar ABC transporter permease | -1.2668 | 3.60E-04 | |
| SMU_1571 | ugpC | ABC transporter ATP-binding protein | -1.1911 | 7.46E-04 | |
| SMU_1590 | | alpha-amylase | -1.4185 | 2.52E-04 | Yes |
| SMU_1591 | ccpA | catabolite control protein A | -1.4661 | 4.63E-04 | Yes |
| SMU_1595 | | carbonic anhydrase | -1.3908 | 2.04E-08 | |
| SMU_1734 | | acetyl-CoA carboxylase carboxyl transferase subunit alpha | -1.8238 | 1.25E-09 | Yes |
| SMU_1735 | accD | acetyl-CoA carboxylase, carboxyltransferase subunit beta | -2.1461 | 8.43E-09 | Yes |
| SMU_1736 | accC | acetyl-CoA carboxylase biotin carboxylase subunit | -2.0359 | 8.40E-10 | Yes |
| SMU_1737 | fabZ | 3-hydroxyacyl-ACP dehydratase FabZ | -1.9017 | 1.10E-09 | Yes |
| SMU_1738 | accB | acetyl-CoA carboxylase biotin carboxyl carrier protein | -2.0349 | 2.29E-10 | Yes |
| SMU_1739 | fabF | beta-ketoacyl-ACP synthase II | -1.9858 | 1.22E-09 | Yes |
| SMU_1740 | fabG | 3-oxoacyl-[acyl-carrier-protein] reductase | -1.9597 | 4.77E-10 | Yes |
| SMU_1741 | fabD | ACP S-malonyltransferase | -1.8476 | 1.00E-09 | Yes |
| SMU_1742c | fabK | enoyl-[acyl-carrier-protein] reductase FabK | -1.7209 | 1.71E-09 | Yes |

| | | | | | |
|---|---|---|---|---|---|
| SMU_1743 | | acyl carrier protein | -1.0489 | 1.01E-06 | |
| SMU_1744 | | beta-ketoacyl-ACP synthase III | -1.8073 | 3.91E-08 | |
| SMU_1745c | | MarR family winged helix-turn-helix transcriptional regulator | -1.9359 | 1.32E-08 | |
| SMU_1812 | | ISL3-like element ISSmu2 family transposase | -1.0243 | 1.14E-03 | |
| SMU_1841 | | sucrose-specific PTS transporter subunit IIBC | -2.4634 | 3.16E-05 | Yes |
| SMU_1843 | | sucrose-6-phosphate hydrolase | -1.6404 | 1.35E-04 | Yes |
| SMU_1844 | | LacI family DNA-binding transcriptional regulator | -1.4908 | 5.83E-04 | Yes |
| SMU_1856c | | TraX family protein | -1.2719 | 4.10E-05 | |
| SMU_1877 | manX | PTS sugar transporter subunit IIB | -2.2329 | 1.62E-04 | Yes |
| SMU_1878 | | PTS mannose/fructose/sorbose transporter subunit IIC | -2.0374 | 1.92E-04 | Yes |
| SMU_1879 | | PTS system mannose/fructose/sorbose family transporter subunit IID | -2.0353 | 8.89E-05 | Yes |
| SMU_1895c | | Blp family class II bacteriocin | -1.5410 | 8.09E-06 | |
| SMU_1924 | gcrR | response regulator GcrR | -1.4267 | 7.48E-07 | |
| SMU_1927 | | ABC transporter ATP-binding protein | -2.0206 | 1.72E-06 | Yes |
| SMU_1928 | | ABC transporter permease; FtsX-like permease family protein | -2.5395 | 4.73E-07 | Yes |
| SMU_1956c | | hypothetical protein | -4.7765 | 2.03E-05 | |
| SMU_1957 | | PTS system mannose/fructose/sorbose family transporter subunit IID | -5.0019 | 1.07E-05 | |
| SMU_1958c | | PTS sugar transporter subunit IIC | -5.0638 | 7.44E-06 | |
| SMU_1960c | | PTS sugar transporter subunit IIB | -5.4203 | 2.21E-06 | |
| SMU_1961c | | PTS sugar transporter subunit IIA | -5.1307 | 8.48E-06 | |
| SMU_2000 | rplQ | 50S ribosomal protein L17 | -1.6450 | 5.94E-10 | |
| SMU_2001 | | DNA-directed RNA polymerase subunit alpha | -1.6056 | 2.24E-09 | |
| SMU_2002 | rpsK | 30S ribosomal protein S11 | -1.5641 | 1.15E-09 | |
| SMU_2003 | rpsM | 30S ribosomal protein S13 | -1.3464 | 8.25E-09 | |
| SMU_2004 | infA | translation initiation factor IF-1 | -1.5022 | 9.25E-09 | |
| SMU_2005 | | adenylate kinase | -1.6492 | 4.65E-10 | |
| SMU_2006 | secY | preprotein translocase subunit SecY | -1.4563 | 2.74E-08 | |
| SMU_2007 | rplO | 50S ribosomal protein L15 | -1.4907 | 2.45E-08 | |
| SMU_2008 | rpmD | 50S ribosomal protein L30 | -1.6160 | 4.96E-09 | |

| | | | | | |
|---|---|---|---|---|---|
| SMU_2009 | rpsE | 30S ribosomal protein S5 | -1.5092 | 8.02E-08 | |
| SMU_2010 | rplR | 50S ribosomal protein L18 | -1.6269 | 6.53E-09 | |
| SMU_2011 | rplF | 50S ribosomal protein L6 | -1.5695 | 9.15E-09 | |
| SMU_2012 | rpsH | 30S ribosomal protein S8 | -1.4847 | 1.31E-08 | |
| SMU_2014 | | type Z 30S ribosomal protein S14 | -1.5785 | 1.14E-08 | |
| SMU_2015 | rplE | 50S ribosomal protein L5 | -1.6522 | 3.76E-09 | |
| SMU_2016 | rplX | 50S ribosomal protein L24 | -1.5856 | 1.12E-08 | |
| SMU_2017 | rplN | 50S ribosomal protein L14 | -1.6980 | 1.75E-08 | |
| SMU_2018 | rpsQ | 30S ribosomal protein S17 | -1.6401 | 6.96E-09 | |
| SMU_2019 | rpmC | 50S ribosomal protein L29 | -1.8014 | 1.13E-08 | |
| SMU_2020 | rplP | 50S ribosomal protein L16 | -1.5926 | 2.61E-09 | |
| SMU_2021 | rpsC | 30S ribosomal protein S3 | -1.6299 | 9.14E-09 | |
| SMU_2022 | rplV | 50S ribosomal protein L22 | -1.6097 | 6.02E-09 | |
| SMU_2023c | rpsS | 30S ribosomal protein S19 | -1.6463 | 4.84E-09 | |
| SMU_2167 | rplB | 50S ribosomal protein L2 | -1.6137 | 1.64E-08 | |
| SMU_2166 | | 50S ribosomal protein L23 | -1.7209 | 4.32E-09 | |
| SMU_2024c | rplD | 50S ribosomal protein L4 | -1.7590 | 9.77E-09 | |
| SMU_2025 | rplC | 50S ribosomal protein L3 | -1.7156 | 4.15E-09 | |
| | rpsJ | 30S ribosomal protein S10 | -1.4787 | 2.20E-09 | |
| SMU_2028 | | glycoside hydrolase family 68 protein | -2.9033 | 5.75E-06 | Yes |
| SMU_2032 | rpsB | 30S ribosomal protein S2 | -1.0659 | 1.01E-06 | |
| SMU_2046c | | endonuclease/exonuclease/phosphatase family protein | -1.4498 | 5.95E-05 | |
| SMU_2047 | | PTS transporter subunit IIBC | -1.7082 | 3.50E-06 | |
| SMU_2080 | brsR | bacteriocin genes transcriptional regulator BrsR; LytTR family transcriptional regulator | -1.0768 | 2.39E-03 | |
| SMU_2127 | | NAD-dependent succinate-semialdehyde dehydrogenase | -2.2005 | 2.25E-03 | |
| SMU_2128 | ilvD | dihydroxy-acid dehydratase | -1.1960 | 4.25E-05 | |

1. Ganguly T, Peterson AM, Burkholder M, Kajfasz JK, Abranches J, Lemos JA. 2022. ZccE is a Novel P-type ATPase That Protects Streptococcus mutans Against Zinc Intoxication. PLoS Pathog 18:e1010477.
